# Rapid bacterial and fungal successional dynamics in first year after Chaparral wildfire

**DOI:** 10.1101/2021.12.07.471678

**Authors:** M. Fabiola Pulido-Chavez, James W. J. Randolph, Cassandra Zalman, Loralee Larios, Peter M. Homyak, Sydney I. Glassman

## Abstract

The rise in wildfire frequency and severity across the globe has increased interest in secondary succession. However, despite the role of soil microbial communities in controlling biogeochemical cycling and their role in the regeneration of post-fire vegetation, the lack of measurements immediately post-fire and at high temporal resolution has limited understanding of microbial secondary succession. To fill this knowledge gap, we sampled soils at 17, 25, 34, 67, 95, 131, 187, 286, and 376 days after a southern California wildfire in fire-adapted chaparral shrublands. We assessed bacterial and fungal biomass with qPCR of 16S and 18S and richness and composition with Illumina MiSeq sequencing of 16S and ITS2 amplicons. Fire severely reduced bacterial biomass by 47%, bacterial richness by 46%, fungal biomass by 86%, and fungal richness by 68%. The burned bacterial and fungal communities experienced rapid succession, with 5-6 compositional turnover periods. Analogous to plants, turnover was driven by “fire-loving” pyrophilous microbes, many of which have been previously found in forests worldwide and changed markedly in abundance over time. Fungal secondary succession was initiated by the Basidiomycete yeast *Geminibasidium*, which traded off against the filamentous Ascomycetes *Pyronema*, *Aspergillus*, and *Penicillium*. For bacteria, the Proteobacteria *Massilia* dominated all year, but the Firmicute *Bacillus* and Proteobacteria *Noviherbaspirillum* increased in abundance over time. Our high-resolution temporal sampling allowed us to capture post-fire microbial secondary successional dynamics and suggest that putative tradeoffs in thermotolerance, colonization, and competition among dominant pyrophilous microbes control microbial succession with possible implications for ecosystem function.

## 1. Introduction

The rapid increase in wildfire frequency, severity, and extent in the western United States (Riley & Loehman, 2016) and around the globe (Abatzoglou et al., 2019) has renewed interest in secondary succession. Secondary succession, or the trajectory along which an ecosystem develops following a disturbance, such as a wildfire, has been extensively studied for plants (Derroire et al., 2016; Donato et al., 2012), but belowground microbial communities have been comparatively overlooked. Understanding how wildfires alter soil microbial succession may be necessary to predict post-fire effects on ecosystem recovery, and function since soil microbes drive post-fire organic matter decomposition (Semenova-Nelsen et al., 2019), nutrient cycling (Pérez-Valera et al., 2020), and plant regeneration (Dove & Hart, 2017).

Plant succession is a major process affecting the health and function of ecosystems. During succession, the dominant species may change in an orderly and predictable manner (Shugart, 2013). For example, early colonizers (fast-growing, short-lived species) are often replaced by late colonizers (slow-growing, long-lived species). However, wildfire can reset the successional clock, shifting the course away from stability (Reilly & Spies, 2016) and initiating secondary succession. Plant secondary succession is often contingent on the surviving vegetation and seedbanks present in the heterogeneous post-fire landscape (Jain et al., 2008). While early plant establishment often happens in low competition and nutrient-rich environments (Dalling, 2008), succession is often mediated by the quantity and identity of early colonizers (i.e., dispersal limitations and priority effects (Kennedy et al., 2009)), and their tradeoffs for space and resources (Tilman, 1990). Indeed, the trajectory of species replacement suggests that early colonizers with similar resource use (i.e., overlapping niches) will dominate open space but will inevitably be replaced by late-stage species with differentiation in resource use (i.e., complementarity), allowing for species coexistence (Pacala et al., 1996; Turnbull et al., 2013). This later stage community is often considered stable, characterized by small fluctuations in community composition. However, whether the patterns recognized in plant successional theory translate to belowground soil microbes remains unclear.

Research on post-fire soil microbiomes suggests that fires can reset microbial successional trajectories via fire-induced mortality (Hart et al., 2005), changes in microbial richness and biomass (Dooley & Treseder, 2012; Pressler et al., 2019), and the replacement of fungal basidiomycetes and symbiotic mycorrhizal fungi with ascomycetes and saprobic fungi (Cairney & Bastias, 2007; Fox et al., 2022). For over a century, pyrophilous or “fire-loving” fungi have been consistently found in post-fire mushroom surveys (McMullan-Fisher et al., 2011; Seaver, 1909). More recently, next-generation sequencing indicates that the pyrophilous Ascomycete *Pyronema* can increase 100-fold after prescribed fires (Reazin et al., 2016) and dominate over 60% of the sequences in experimental pyrocosms (Bruns et al., 2020). Furthermore, there is increasing evidence of pyrophilous bacteria, such as the Proteobacteria *Massilia* (Enright et al., 2022; Whitman et al., 2019). This evidence suggests that pyrophilous microbes in systems that have evolved with wildfire (Dove et al., 2021) might have fire adaptations analogous to plants (Rundel, 2018) and, thus, likely follow successional dynamics akin to plants. Pyrophilous microbes have traits that allow them to survive fires (e.g., heat resistant spores, sclerotia) (Day et al., 2020; Petersen, 1970) and the post-fire environment (e.g., xerotolerance, affinity for nitrogen mineralization, and affinity for aromatic hydrocarbon degradation) (Fischer et al., 2021; Nelson et al., 2022; Steindorff et al., 2021). Moreover, since heat from fire often penetrates only the top few cm of soil (Neary et al., 1999; Pingree & Kobziar, 2019), like plants, secondary succession may be initiated by surviving microbes that make up the spore bank (Baar et al., 1999; Glassman et al., 2016). Currently, most research evaluating post-fire microbiomes is based on single timepoint sampling (Dove & Hart, 2017; Pressler et al., 2019); thus, the succession of pyrophilous microbes is nearly unknown. However, recent research consisting of 2-3 sampling time points suggests that bacteria and fungi experience rapid post-fire community changes (Ferrenberg et al., 2013; Qin & Liu, 2021; Whitman et al., 2022), indicating that higher temporal resolution sampling is needed to understand bacterial and fungal successional trajectories.

Predicting soil microbial succession can be complicated by direct and indirect wildfire effects (Neary et al., 1999). Whereas soil burn severity controls direct fire impacts (Reazin et al., 2016; Whitman et al., 2019), changes in soil moisture can indirectly impact post-fire microbes (Placella et al., 2012; Yang et al., 2021), potentially driving succession, especially in arid environments where precipitation is limited. Previous research has established that bacteria rapidly respond to soil wet-up (Barnard et al., 2013; Placella et al., 2012), whereas fungi are less responsive to soil moisture changes (Barnard et al., 2013; Evans & Wallenstein, 2012). Furthermore, microbial life strategies can determine microbial response to fire. Research shows that fire decreases the richness of ectomycorrhizal fungi (EMF) (Cowan et al., 2016; Glassman et al., 2016; Pulido-Chavez et al., 2021) and arbuscular fungi (AMF) (Xiang et al., 2015) but can temporarily increase saprobic richness (Enright et al., 2022; Semenova-Nelsen et al., 2019). Yet, the limited studies on how fires affect multiple microbial guilds (Certini et al 2021) lack the resolution to identify the direct effect of wildfire and the length of time that mycorrhizal fungi survive after a wildfire, a question essential for post-fire ecosystem recovery.

We propose that microbial ecological succession theory can be developed and tested in nature by focusing on California chaparral. Chaparral is a shrubland adapted to high-severity fire and a biodiversity hotspot distributed in Mediterranean climates worldwide (Barro & Conard, 1991; Rundel, 2018). Chaparral plant secondary succession is relatively well understood and typically initiated by fire (Keeley et al., 2005). Yet, although the dominant chaparral vegetation forms mycorrhizal associations required for their establishment and survival (Allen et al., 2005), little is known about chaparral microbial succession. Indeed, only 13% of post-fire microbiome research occurs in shrublands (Pressler et al., 2019). With drylands covering nearly 41% of Earth’s land surface and expanding with climate change (Feng & Fu, 2013), understanding secondary successional dynamics in dryland systems such as chaparral is critical (Osborne et al., 2022). While our focus is on chaparral, which is likely to have fire-adapted microbes due to chaparrals’ evolutionary history with fire (Hanes, 1971), we expect that successional patterns will be broadly generalizable to other biomes since pyrophilous bacteria and fungi appear to be phylogenetically conserved (Enright et al., 2022). Indeed, fungal taxa within the Ascomycota family Pyronemataceae and a few Basidiomycota genera appear to be globally distributed in fire-disturbed Spanish shrublands (Pérez-Valera et al., 2018) and Pinaceae (Reazin et al., 2016; Whitman et al., 2019; Xiang et al., 2014), Eucalyptus (Ammitzboll et al., 2022; McMullan-Fisher et al., 2011), and redwood-tanoak forests (Enright et al., 2022).

Here, we performed the highest resolution temporal sampling of post-fire microbiomes to date. We sampled soils at nine timepoints over one year following a chaparral wildfire, allowing us to identify immediate and temporal effects of wildfire on microbial successional dynamics and test the following hypotheses: (H1) wildfire will decrease bacterial and fungal biomass and richness, leading to a shift in the community composition one year post-fire; (H2) wildfire will have a distinct impact on fungal guilds with symbiotic mycorrhizal fungi experiencing the largest declines; (H3) higher soil burn severity will lead to larger reductions in both bacteria and fungi while precipitation will impose a larger impact on bacterial than fungal succession; (H4) succession will be initiated by pyrophilous microbes, which like plants, will change in abundance over time based on differences in ecophysiological traits such as growth and nutrient acquisition.

## 2. Methods

### 2.1 Study Area, plot design, and soil collection

The Holy Fire burned 94 km^2^ in the Cleveland National Forest in Southern California from August 6 to September 13, 2018. On September 30, 2018, we selected nine plots (6 burned and 3 unburned) (Fig. 1A). Plots were selected for similarity in aspect, slope, elevation, and pre-fire vegetation dominance by manzanita (Arctostaphylos glandulosa), an ectomycorrhizal host, and chamise (Adenostoma fasciculatum), an arbuscular and ectomycorrhizal host (Allen et al., 2005). Plots were placed on average 25 m from forest access roads (10-40 m) to avoid edge effects, and each contained four 1 m^2^ subplots located 5 m from the center in each cardinal direction (Fig. 1B). Our study site experiences a Mediterranean-type climate with hot, dry summers and cool, wet winters, with an average yearly temperature of 17°C and total precipitation of 668 mm (average 51.39 mm). Precipitation data, based on monthly summaries, was gathered from the El Cariso weather station (raws.dri.edu). Soils are mapped in the Cieneba and Friant series and are classified as Typic Xerorthents and Lithic Haploxerolls. They are sandy and gravelly loams with an average pH of 6.8 for the burned plots and 6.2 for the unburned plots (additional plot information in Table S1). Although soil types differ, soil type did not affect bacterial or fungal biomass or richness (glmer.nb; p > 0.05).

**Figure 1.**
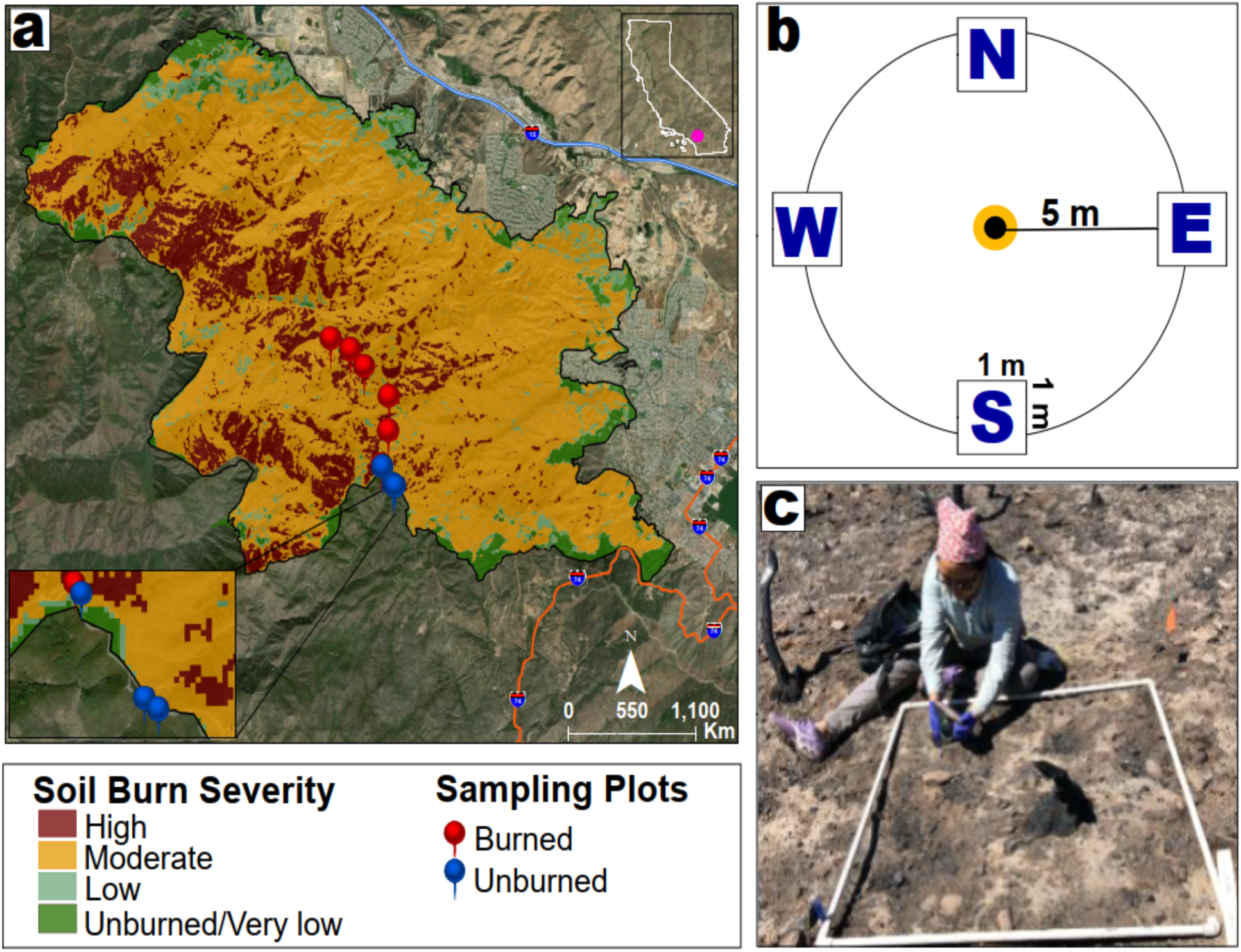
A) Location of the study area and plots (6 burned; 3 unburned) within the Holy Fire burn scar in the Cleveland National Forest in southern California. Soil burn severity is based on the USDA BAER classifications made at seven days within fire containment. B) Plot experimental design, each with four 1m^2^ subplots placed 5m from the center in each cardinal direction. C) Collection of the top 10 cm of soil beneath the ash with a releasable bulb planter within the 1 m^2^ subplots at 17 days post-fire.

We used BAER soil burn severity maps created within seven days after fire containment (https://burnseverity.cr.usgs.gov) to locate plots within moderate burn severity (Fig. 1A). Notably, the BAER coarse-scale measurement (30 m) does not coincide well with the spatial distribution and turnover rate of soil microbial communities (Bahram et al., 2013). Since ash depth increases with soil heating (Parson et al., 2010) and fire severity (Bodí et al., 2014), we used ash depth as a proxy for soil burn severity by averaging three separate measurements of ash depth (cm) per each 1 m^2^ subplot (Fig. 1C). Since ash can be rapidly redistributed via post-fire wind and rain (Bodí et al., 2014), we measured ash before any precipitation event (first rains occurred on Oct 3) or high wind events (average wind Sept 3 -Sept 17 was 2.7 m/s). Hence, our measure of soil burn severity refers to initial ash depth.

We sampled soils at 17, 25, 34, 67, 95, 131, 187, 286, and 376 days, corresponding to approximately 2 and 3 weeks, and 1, 2, 3, 4, 6, 9, and 12 months post-fire. At each of the nine time points, we collected the top 10 cm of mineral soil (A horizons) beneath the ash layer from each burned subplot. Since wildfire results in the combustion of the organic layer, to ensure homogeneity in sampling and direct comparison between treatments in the unburned plots, we removed the litter layer before sampling A horizons (Pulido-Chavez et al., 2021). Soils were collected with a ∼250 mL releasable bulb planter cleaned with ethanol after each use to prevent cross-contamination, resulting in 36 soil samples (9 plots x 4 subplots) per sampling time point. Soils were transported in individual Whirl-Paks in a cooler to the University of California-Riverside (UCR) within hours of sampling, stored overnight at 4°C, and sieved (2mm) at room temperature in ethanol cleaned sieves within 24 hours of sampling to minimize microbial community turnover (Phillips, 2021). A subsample was frozen at -80°C for future DNA extraction. While soils were stored at -80°C from 1 month to 1 year before DNA extractions were performed, differences in storage time at -80°C do not affect DNA quantity and integrity (Lauber et al., 2010; Pavlovska et al., 2021).

### 2.3 DNA extraction, amplification, and sequencing

DNA extractions for all nine timepoints were performed in the summer of 2019. Soils were weighed (0.25 g) using ethanol-cleaned spatulas and processed using Qiagen DNeasy PowerSoil Kits following the manufacturer’s protocol, with an increase in centrifugation time to 1.5 min after adding the C3 solution (a solution used to precipitate the organic and inorganic matter) due to large amounts of precipitate still in the solution, then stored at -20°C. Extracted DNA was amplified using the primer pair ITS4-fun and 5.8s to amplify the ITS2 region for fungi (Taylor et al., 2016) and the primer pair 515F-806R to amplify the V4 region of the 16S rRNA gene for archaea and bacteria (Caporaso et al., 2011) using the Dual-Index Sequencing Strategy (DIP) (Kozich et al., 2013). Although our 16S primers amplify archaea and bacteria, for simplicity, we refer to 16S methods and results simply as bacteria since archaea only contributed <1% of sequencing reads. We conducted polymerase chain reaction (PCR) in two steps. The first PCR amplified gene-specific primers, and the second PCR ligated the DIP barcodes and adaptors for Illumina sequencing. For bacteria, we combined 1 μL of 1:10 diluted DNA, 10.5 μL of Ultra-Pure Sterile Molecular Biology Grade water (Genesee Scientific, San Diego, CA, USA), 12.5 μL of AccuStart ToughMix (2x concentration; Quantabio, Beverly, MA, USA), and 0.5 μL each of the 10 μM 515F and 806R primers. Thermocycler conditions for PCR1 were: 94°C for 2 min followed by 29 cycles of 94°C for 30 s, 55°C for 30 s, 68°C for 1 min, followed by a 2 min extension at 68°C. For fungi, we combined 5 μL of undiluted DNA, 6.5 μL of ultra-pure water, 12.5 μL of AccuStart ToughMix, and 0.5 μL each of the 10 μM ITS4-fun and 5.8s primers. Thermocycler conditions for PCR1 were: 94 °C for 2 min, followed by 30 cycles of 94 °C for 30 s, 55 °C for 30 s, 68 °C for 2 min, followed by a 10 min extension at 68°C. PCR1 products were cleaned with AMPure XP magnetic beads (Beckman Coulter Inc., Brea, CA), following the manufacturer’s protocols. The DIP PCR2 primers containing the barcodes and adaptors for Illumina sequencing were ligated to the amplicons during the second PCR step in a 25 μL reaction containing 2.5 μL of the 10 μM DIP PCR2 primers, 6.5 μL of ultra-pure water, 12.5 μL of Accustart ToughMix, and 1 μL of PCR1 product. Thermocycler conditions for PCR2 for bacteria and fungi were: 94°C for 2 min followed by 10 cycles of 94°C for 30 s, 60°C for 30 s, and 72°C for 1 min. Bacterial and fungal PCR products for the 324 samples (9 plots x 4 subplots x 9 timepoints) were then separately pooled based on gel electrophoresis band strength and cleaned with AMPure following established methods (Glassman et al., 2018). Each pool contained 2-3 timepoints from burned and unburned plots and was checked for quality and quantity with the Agilent Bioanalyzer 2100 before combining bacteria and fungi at a 2:3 ratio (0.4 units for bacteria to 0.6 units for fungi). Since each library only fit 2-3 timepoints for both bacteria and fungi, we sequenced the nine timepoints across four libraries with Illumina MiSeq 2×300bp at the UCR Institute for Integrative Genome Biology. In addition to our 324 experimental samples, we added negative DNA extractions and PCR controls from each time point and mock communities (ZymoBIOMICS microbial community standard, Zymo, Irvine, CA) to each library for additional quality control and inference.

### 2.4 Bacterial and fungal biomass

We estimated bacterial and fungal gene copy numbers with quantitative (q) PCR as a proxy for biomass using the Eub338/Eub518 primers for bacteria (Fierer et al., 2005) and FungiQuant-F/FungiQuant-R primers for fungi (Liu et al., 2012). We used the small subunit since it is more conserved and has less length variation making it better suited for qPCR than the ITS2 region (Mayer et al., 2021), which is better at species level identification (Schoch et al., 2012). We generated standard curves using a 10-fold serial dilution of the standards by cloning the 18S region of the fungus *Saccharomyces cerevisiae* or the 16S region of the bacteria *Escherichia coli* into puc57 plasmid vectors constructed by GENEWIZ, Inc. (NJ, USA) as previously established (Averill & Hawkes, 2016). The 10 μL qPCR reactions were performed in triplicate. Reactions contained 1 μL undiluted DNA, 1 μL of 0.05M Tris-HCl ph8.3, 1 μL of 2.5mM MgCl_2_ (New England BioLabs (NEB); Ipswich, MA), 0.5 μL of 0.5mg/ml BSA, 0.5 μL of 0.25mM dNTP (NEB), 0.4 μL of both primers at 0.4μM, 0.5 μL of 20X Evagreen Dye (VWR International; Radnor, PA), 0.1 μL of Taq DNA polymerase (NEB) and 4.6 μL ultra-pure water. We employed the CFX384 Touch Real-Time PCR Detection System with the following conditions: 94°C for 5 min, followed by 40 cycles of 94°C for 20 seconds, 52°C (for bacteria) or 50°C (for fungi) for 30 seconds, followed by an extension at 72°C for 30 seconds. Gene copy numbers were generated using the equation 10^^^ (Cq-b)/m), using the quantification cycle (Cq), calculated as the average Cq value per sample in relation to the known/calculated copies in the Cq/threshold cycles, the y-intercept (b) and the slope (m) generated with CFX Maestro software. Values with an R^2^ > 0.994 were considered acceptable. Gene copy numbers were normalized per gram of dry soil (Tatti et al., 2016). Notably, all methods of estimating microbial biomass have limitations (Gao et al., 2022), including qPCR, due to variation in target copy number (Lofgren et al., 2019; Song et al., 2014). However, the high sensitivity (Tellenbach et al., 2010) of qPCR for quantifying the selected marker gene (Smith & Osborn, 2009) makes it ideal for estimating microbial biomass from environmental samples when interest is not in species specificity but in estimates of total microbial biomass. Moreover, common biomass quantification methods for fungi are correlated (Cheeke et al., 2017).

### 2.5 Bioinformatics

Illumina data was processed with Qiime2 version 2020.8 (Bolyen et al., 2019). The demultiplexed fastQ files from the four Illumina sequencing runs were processed individually using cutadapt (Martin, 2011) to remove the primers and ran through DADA2 version 2020.8 with the defaults parameters to filter out and remove chimeric sequences and low-quality regions and to produce Amplicon Sequence Variants (ASVs) (Callahan et al., 2017). Bacteria forward reads were trimmed to 170 bp and reverse to 163 bp, while fungal forwards reads were trimmed to 209 bp and the reverse reads to 201 bp. On average, 72% of forward and reverse sequences merged for bacteria, and 61% merged for fungi. DADA2 outputs from each library were combined into one library for downstream processing, including the removal of singletons and taxonomic assignments. We used SILVA version 132 for bacterial (Yilmaz et al., 2014) and UNITE version 8.2 for fungal (Abarenkov et al., 2020) taxonomic assignment using Qiime2 Naïve Bayes Blast+ classifier. Bacterial sequences assigned to mitochondria and chloroplasts and fungal sequences not assigned to Kingdom Fungi were removed. Fungal ASV tables were exported and parsed through FUNGuild (Nguyen et al., 2016) to assign functional ecological guilds, including only highly probable confidence rankings for AMF, EMF, saprotrophs, and pathogens. Moreover, negative and mock controls were inspected to ensure that sequences and/or ASVs in negative controls were negligible and that mock taxonomy correlated with known communities. Sequences were submitted to the National Center for Biotechnology Information Sequence Read Archive under BioProject accession number PRJNA761539.

### 2.6 Statistical analysis

The ASV tables were rarefied to a sequences/sample depth of 7,115 for bacteria and 11,480 for fungi to account for uneven sequencing depth. Fungal and bacterial alpha diversity was estimated using BiodiversityR version 2.14-2 (Kindt & Coe, 2005) with the following metrics: observed species richness, Simpson, Shannon, Chao1, ACE, Simpson’s evenness, and Inverse Shannon. Although estimating richness from diverse soil communities has limitations (Willis et al., 2017), wildfire-affected soil constitutes low diversity communities, and patterns of species richness were similar across metrics (Fig. S1). Thus, we focused on richness estimated as the number of observed ASVs after rarefaction for all downstream analyses. To test if wildfire decreased biomass and richness and if biomass and richness increased with time since fire (H1), if wildfire will have a distinct impact on fungal guilds (H2), and to determine if changes in biomass and richness are associated with precipitation and soil burn severity (H3), for both bacteria and fungi, we performed backward model selection and fitted nine statistical models with treatment (burned vs. unburned), time (measured as days since fire), ash depth (cm), total monthly precipitation (mm), and second order interactions as predictors. We used generalized mixed effect models (glmer) with a negative binomial distribution in the MASS package version 7.3-57 (Venables & Ripley, 2002) to account for the over-dispersion of the data and the fact that the conditional variance was higher than the conditional mean (Bliss, 1953; Ross & Preece, 1985). Time since fire and precipitation were scaled and centered. The level of nestedness for all models was tested by running a null model with different nested levels (plot, subplot, and time since fire) and no predictors. Model selection was made using Akaike Information Criterion (AICc) in the MuMIN package version 1.46.0 (Barton, 2020). All richness models contained plot, subplot, and time since fire as random effects, and all biomass models included plot and time since fire as random effects. The subplot was not included for biomass analysis as it was not determined to be the best model (AIC selection). Pseudo R^2^, or the variance explained (marginal and conditional) for all models, was calculated using the r.squaredGLMM function in the MuMIn Package.

We compared beta diversity by generating distance matrices with the vegan Avgdist function, calculating the square-root transformed median Bray-Curtis dissimilarity per 100 iterations. We used permutational multivariate analysis of variance (PERMANOVA) (Anderson, 2017) as implemented with the adonis function in vegan version 2.6-2 (Oksanen et al., 2018) to test for the significant effects of wildfire, time since fire, precipitation, ash depth, and second order interactions, on bacterial and fungal community composition overall. Moreover, we tested the significance of wildfire at each independent time point (H1). Results were visualized using Non-Metric Multidimensional Scaling (NMDS) ordinations.

We employed several methods to quantify succession (H4). First, we visualized succession by characterizing community composition patterns at the genus level using phyloseq version 1.38.0 (McMurdie & Holmes, 2013) and grouping the relative abundance of the dominant ASVs (> 3% sequence abundance used for plot legibility) for each time point in burned and unburned communities independently. Second, we used the vegan mantel function to determine the correlation between temporal distance and community composition to determine how turnover varies in the presence or absence of fire, similar to a study examining Sorghum microbial succession (Gao et al., 2020). Third, to visualize spatial species turnover (Baselga, 2010), we took advantage of the fact that early successional periods display large variability (Collins, 1990; Pandolfi, 2008). Thus, we tested the homogeneity of the bacterial and fungal communities during succession using the vegan betadisper function to calculate the multivariate dispersion of Bray-Curtis dissimilarities and visualized the results using principal coordinates analysis (PCoA). Finally, we employed the codyn package version 2.0.5 (Hallett et al., 2016) to identify the patterns of community dynamics over successional time, including the rate of directional change, stability, and synchrony (Collins et al., 2000). Succession involves directional change via the replacement of early to late successional species (Clements, 1916; Platt & Connell, 2003); thus, we calculated the rate of directional change using the Euclidian distance of each microbial community. Then we measured species turnover to determine if species appearances or disappearances drive succession. Since stability increases as diversity increases (diversity-stability hypothesis), we independently measured community stability in the burned and unburned plots as the mean abundance divided by the standard deviation at each temporal time point (Lehman & Tilman, 2000). However, community stability depends on species covariance over time; thus, we measured synchrony (Loreau & de Mazancourt, 2008), where perfect species synchrony is a value of 1 and 0 equals asynchrony (Valencia et al., 2020). Asynchrony can result in covariance in species populations, where tradeoffs among species can contribute to overall community stability.

All statistical analyses were conducted in R 4.1.1 (R Core Team, 2021), and plots were created with ggplot2 version 3.3.6 (Wickham, 2016). All statistical codes are available on GitHub https://github.com/pulidofabs/SecondarySuccession-Chaparral.

## 3. Results

### 3.1 Sequencing data

The four Illumina MiSeq runs resulted in 9.8 M bacterial and 24.6 M fungal sequences for an average of 31,052 bacterial and 78,202 fungal sequences/sample and a total of 33,078 bacterial and 11,480 fungal ASVs. Compared to the experimental samples, bacterial negative controls, on average, had 10 ASVs, while fungal negative controls had 8 ASVs (Table S2). Rarefaction resulted in a total of 24,874 bacterial and 7,445 fungal ASVs, and the removal of all control samples. Extracting the fungal guilds from the rarefied fungal ASV table resulted in 208 EMF, 70 saprobic, 65 AMF, and 26 pathogenic fungal ASVs.

### 3.2 Wildfire effects on bacterial and fungal biomass and richness

Fire significantly reduced bacterial and fungal biomass and richness during the first post-fire year (Tables S3). Fire had a larger effect on biomass than richness for both bacteria and fungi and larger effects on fungi than bacteria overall (Fig. 2). On average, across the year, fire reduced bacterial richness by 46% (Fig. S1A) and biomass by 47% (Fig. S2A) and fungal richness by 68% (Fig. S1A) and biomass by 86% (Fig. S2B). The direct, immediate effect of wildfire at 17 days post-fire was stronger for biomass than richness, with bacterial biomass decreasing by 84% (Fig. 2A) and fungal biomass by 97% (Fig. 2B; Table S3). In contrast, at 17 days post-fire, fungal richness declined by 45% (Fig. 2D), but bacterial richness temporarily increased by 31% but declined by 29% at 25 days post-fire (Fig. 2C; Table S3). While the differences in biomass and richness between the burned and unburned plots lessened with time since fire, one year was insufficient for bacterial and fungal biomass or richness to recover to unburned levels (Fig. 2). The effects of fire across time remained larger for fungi than for bacteria, with fungal biomass remaining 80% and richness 61% lower and bacterial biomass remaining 43% and richness 23% lower in the burned plots at 1-year post-fire (Fig. 2; Table S3).

**Figure 2.**
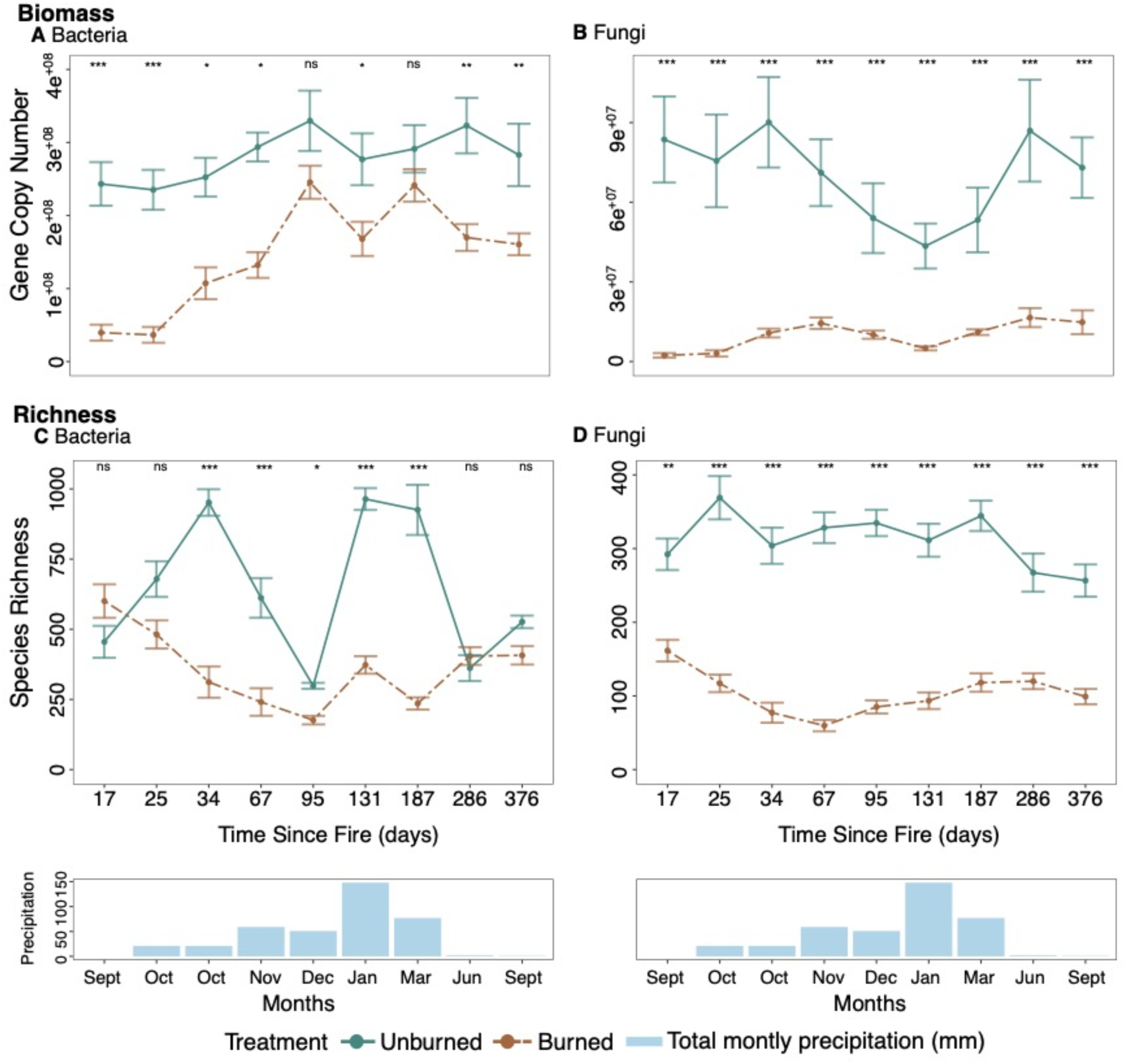
Change between burned (brown) and unburned communities (blue-green) for A) bacterial and B) fungal estimated biomass per gram of soil and C) bacterial and D) fungal species richness at each of the nine timepoints in the first post-fire year. Points represent the mean biomass (A, B) and richness (C, D), and bars represent the standard error of the mean for burned versus unburned plots at each timepoint. Note that gene copy number and species richness are displayed on a daily scale, whereas precipitation is on a monthly scale representing total precipitation for the month in which each soil sample was collected. Note, we sampled twice in Oct. The significance of burned versus unburned plots per timepoint is denoted with an asterisk and is based on a negative binomial regression.

### 3.3 Fire directly and indirectly affects biomass and richness

Fire had direct, negative effects on bacterial and fungal biomass and richness. These direct effects were modified over time by positive interactions with fire for fungal biomass and with soil burn severity for fungal richness and bacterial biomass (Fig. 3; Table S4). Time since fire also directly positively impacted fungal richness (Table S4). Soil burn severity had significant adverse direct effects on bacterial and fungal richness but no significant direct effects on biomass, meaning that biomass was equally reduced by fire regardless of severity (Figs. 3C, 3D; Table S4). While precipitation had no significant direct effects, precipitation interacted with fire to positively impact bacterial biomass (Fig. 2A), and precipitation interacted with time since fire to negatively affect bacterial richness (Fig. 2C) but positively affect fungal richness (Fig. 2D; Table S4).

**Figure 3.**
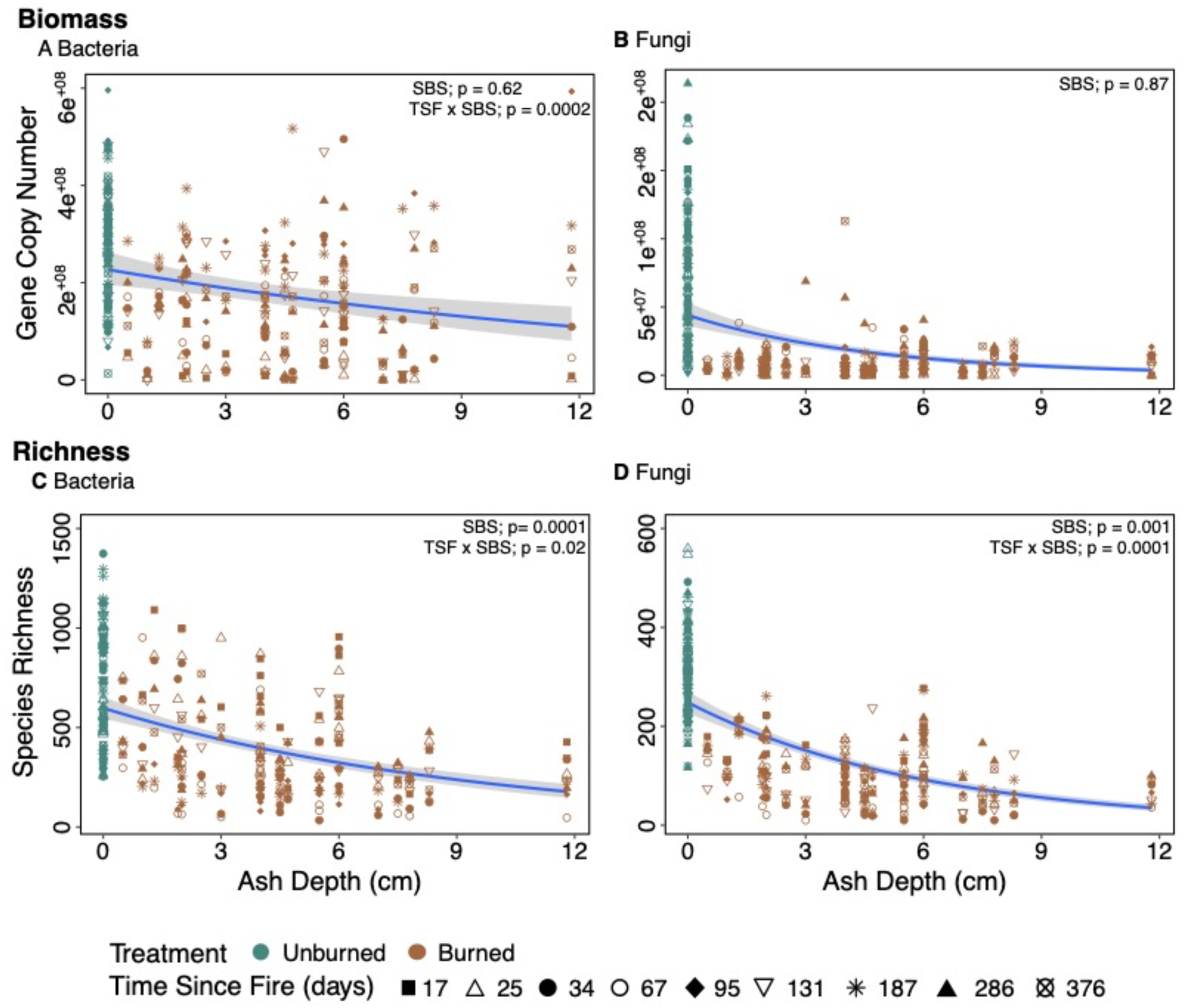
Change in estimated biomass and richness with soil burn severity measured as ash depth at 17 days post-fire for A) bacterial and B) fungal biomass per gram of soil and C) bacterial and D) fungal richness. The significance of biomass and species richness against soil burn severity (SBS) and its interaction with time since (TSF) is based on a negative binomial regression (Table S4). The blue line represents the model’s prediction, and the gray represents the standard error.

### 3.4 Wildfire effects on the richness of different fungal guilds

Wildfire led to large and significant reductions in species richness for all fungal guilds (Fig. S3; Table S5), with the largest initial declines for AMF. On average, across time points, EMF decreased by 68%, pathogens by 71%, saprobes by 86%, and AMF by 98% in burned compared to unburned plots (Fig. S3), and one year was insufficient for any guild to return to unburned levels. In fact, at 376 days, EMF were on average 91%, AMF 89%, pathogens 69%, and saprobes 61% lower in the burned plots (Fig. S4; Table S5). EMF richness declined with soil burn severity and due to a fire and time interaction, whereas time since fire had a direct, negative effect on saprobic richness, which declined over time (Table S5). Finally, an interaction between fire and precipitation negatively affected both EMF and pathogen richness (Table S5).

### 3.5 Wildfire changes microbial community composition

Fire significantly affected bacterial and fungal community structure, explaining 13% of the compositional variation for bacteria and 10% for fungi (Table S6). Time since fire had smaller impacts on community composition, explaining 4% of the variation for bacteria and 1% for fungi. Ash depth equally affected bacterial and fungal composition explaining 2% of the variance for both, whereas precipitation had slightly larger impacts on bacteria than fungi, explaining 3% of bacterial and 1% of the fungal variation (Table S6). There were also small but significant interaction effects for both bacteria and fungi (Table S6). Furthermore, community composition between burned and unburned plots significantly varied at all 9-time points for both bacteria (Fig. S5) and fungi (Fig. S6), with differences in community composition increasing over time from 12% to 21% for bacteria (Fig. S5) and 9% to 13% for fungi (Fig. S6) from day 17 to 376 post-fire. The overall changes in burned bacterial (Fig. 4A) and fungal composition (Fig. 4C) were driven by the previously rare taxa, which increased in abundance over time (Figs. 4B, 4D). Unlike fungi, burned bacterial communities were consistently dominated over time (57% average sequence abundance) by a single genus, the Proteobacteria *Massilia* (Fig. 4B). However, three Firmicutes quickly rose in dominance, with *Bacillus* (13%), an uncultured Clostridiales (26%), and *Paenibacillus* (4%) dominating at 34- and 67-days post-fire. Yet these Firmicutes rapidly declined as the Proteobacteria *Noviherbaspirillum* increased in abundance over time, from 1% at 34 days to 24% by the end of the year (Fig. 4B). For burned fungi, two Basidiomycetes that dominated at 17 days post-fire, the yeast *Geminibasidium* (45%) and the EMF *Inocybe* (16%), rapidly declined while filamentous Ascomycetes increased over time (Fig. 4D). The genera *Pyronema* increased as *Geminibasidium* decreased and dominated from 25 to 95 days post-fire, peaking at 67 days post-fire with 67% sequence abundance. In addition, the Ascomycetes *Aspergillus* increased from 2% to 22% and *Penicillium* from 36% to 49% from 17 to 376 days post-fire (Fig. 4D).

**Figure 4.**
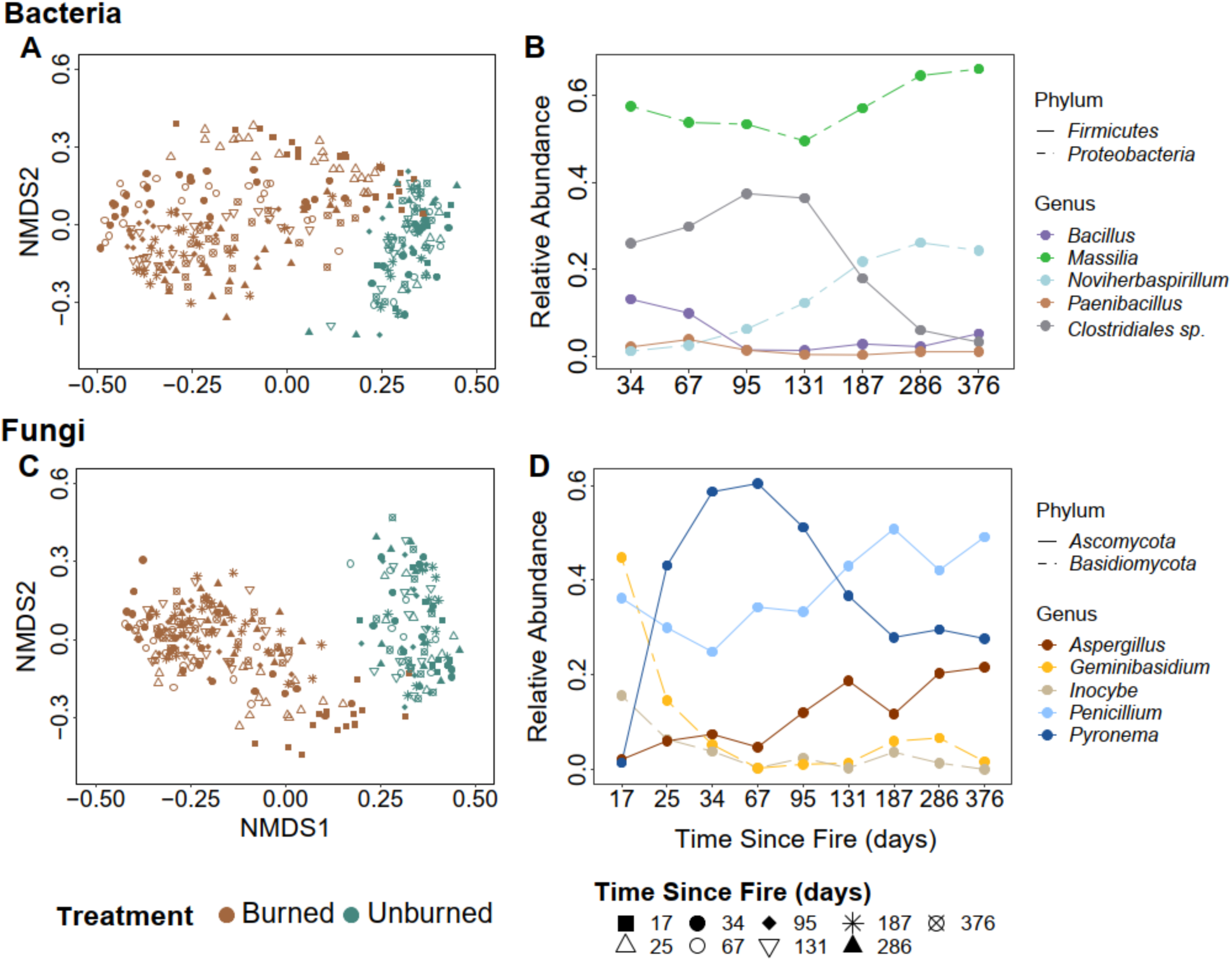
NMDS plot of A) bacterial and C) fungal community composition with colors denoting treatment and shape indicating time since fire. The NMDS is based on 3-dimensions and has a stress value of 0.11 for bacteria and 0.12 for fungi. Change in relative sequence abundance of the five most abundant B) bacterial and D) fungal genera in the burned plots over time. Note that the dominance of the top 5 bacterial genera began 34 days post-fire since there was no clear dominance on days 17 and 25 due to the lag in post-fire bacterial declines.

### 3.6 Bacterial successional dynamics

Burned bacterial communities experienced rapid and distinct successional trajectories (Fig. 5A) at roughly double the rate of fungi (Fig 5C; Fig. 5D) and were driven by six major compositional turnover points at 25, 34, 95, 131, 187, and 376 days (Fig. 5E). In contrast, the unburned bacterial communities experienced low dominance and remained stable over time (Fig. 6A) with no succession (Figs. 6C) or compositional turnover (Figs. 6E). These patterns are also reflected in the measures of successional dynamics (Table S7). For example, burned bacterial communities experienced temporal changes in species composition that were directional and higher (0.16) than in the unburned communities (0.12; Table S7). Burned bacterial communities also exhibited lower synchrony (0.03) than unburned communities (0.22), resulting in higher stability in burned (8.35) compared to unburned bacterial communities (5.36; Table S7).

**Figure 5.**
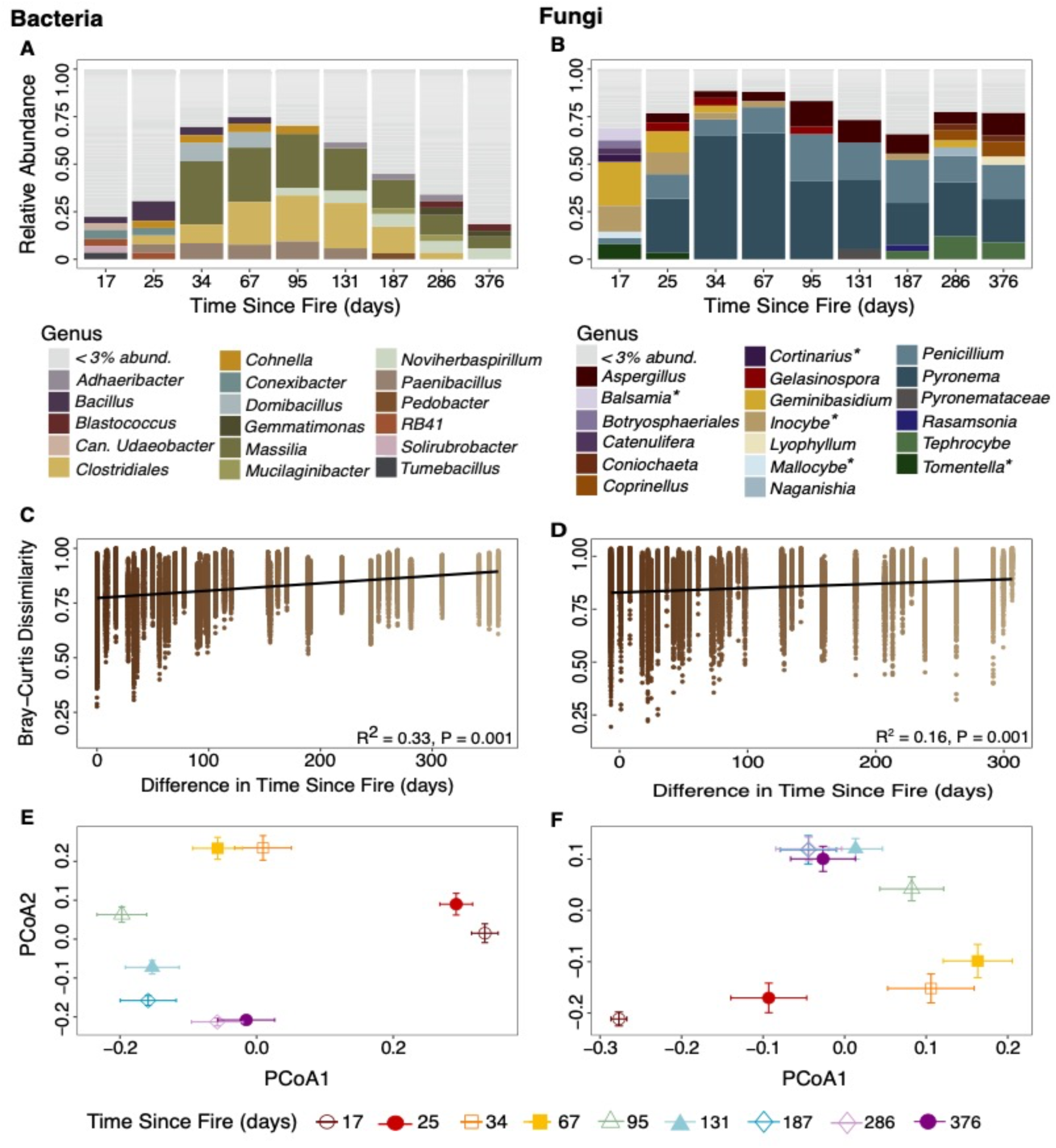
Relative sequence abundance of A) bacterial and B) fungal genera (ectomycorrhizal fungi denoted with an asterisk) in burned plots at all 9 timepoints. Mantel correlation between Bray-Curtis dissimilarity and Euclidean temporal distances in C) bacterial and D) fungal community composition in burned plots. Principal components analysis (PCoA) of the mean and standard error Bray-Curtis community dissimilarity at each timepoint for the burned E) bacterial and F) fungal communities. Note that timepoints farther apart with non-overlapping standard error bars indicate a community turnover, whereas overlapping standard error bars represent a lack of compositional turnover. Note that 17 days represents the base level. Thus, the first turnover was initiated at 25 days. Gray bars represent multiple genera of bacteria (A) and fungi (B) that have relative sequence abundance under 3% at each timepoint per treatment. Note that 47% of the bacterial and 20% of the fungal sequences comprise genera <3% relative abundance.

**Figure 6.**
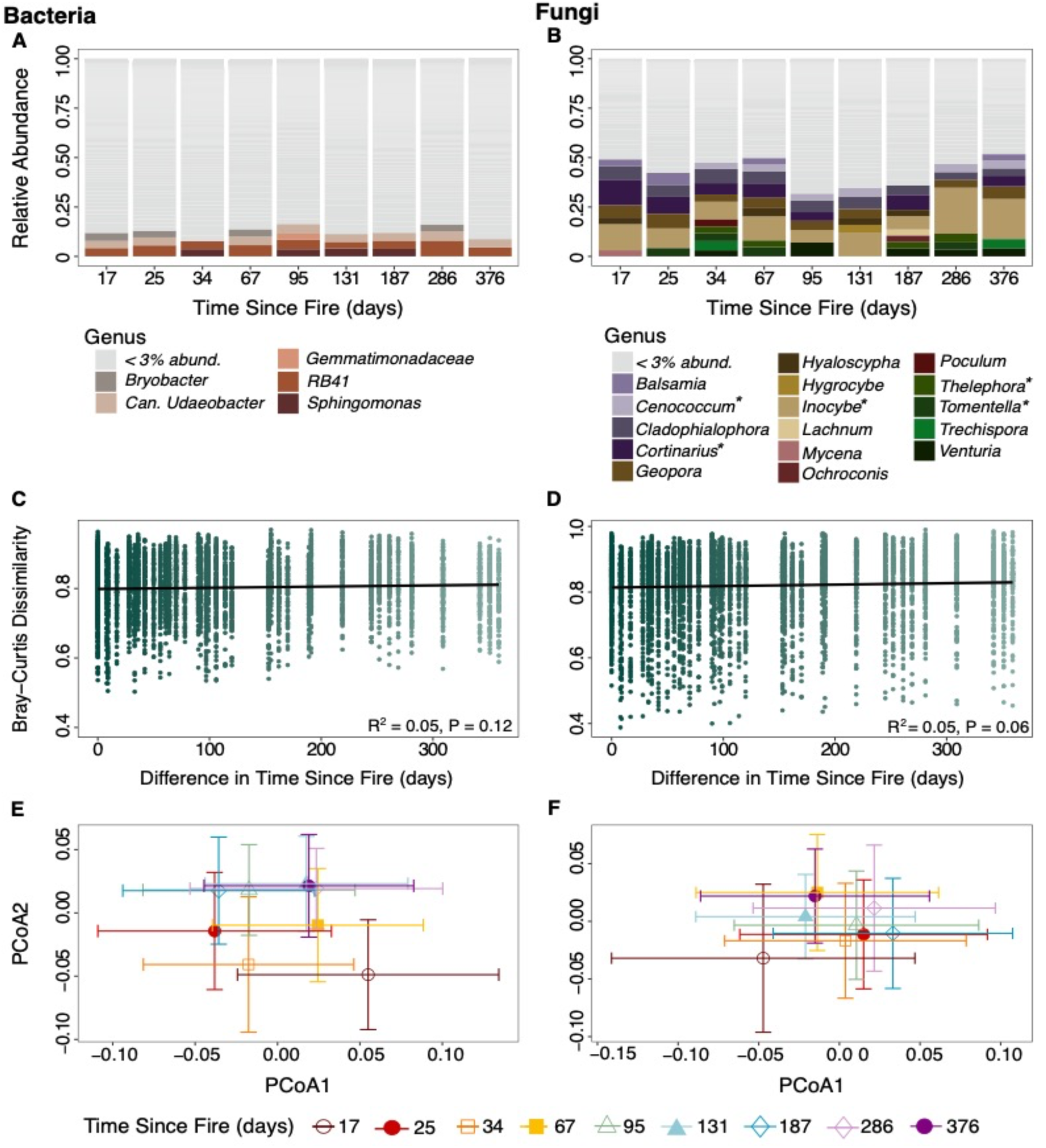
Relative sequence abundance of A) bacterial and B) fungal genera (ectomycorrhizal fungi denoted with an asterisk) in unburned plots at all 9-time points. Mantel correlation between Bray-Curtis dissimilarity and Euclidean temporal distances in C) bacterial and D) fungal community composition in unburned plots displaying no significant succession. Principal components analysis (PCoA) of the mean and standard error Bray-Curtis community dissimilarity at each time point for E) bacteria and F) fungi in the unburned plots. In unburned plots, overlapping standard error bars indicate no community turnover across time. Gray bars represent multiple genera of bacteria (A) and fungi (B) that have relative sequence abundance under 3% at each timepoint per treatment. Note the lack of dominance as denoted by the fact that 79% of the bacterial and 51% of the fungal sequences comprise genera <3% relative abundance.

### 3.7 Fungal successional dynamics

Burned fungal communities experienced rapid and distinct successional trajectories (Fig. 5B; Fig. 5D) that were driven by five major compositional turnover points at 25, 34, 67, 95, and 131 days post-fire (Fig. 5F). In contrast, the unburned fungal communities experienced low dominance (Fig. 6B) and remained stable over time with no succession (Fig. 6D) or compositional turnover (Fig. 6F). Like bacteria, these patterns were similarly reflected in the measures of successional dynamics (Table S7) such that burned fungal communities had higher directional change (0.49) than the unburned communities (0.08), reflecting stronger patterns of succession as more species were introduced into the burned system. Burned fungal communities exhibited slightly lower synchrony (0.04) than unburned communities (0.05) but much lower community stability (6.42) than unburned communities (8.58), indicating higher susceptibility to change in burned fungal communities (Table S7).

### 3.8 Taxa driving microbial succession

We found that bacterial and fungal succession were driven by taxa that traded off in abundance based on physiological traits, such as thermotolerance and fast colonization (Table 1). Early turnover events for bacteria (Fig. 5A) and fungi (Fig. 5B) in burned plots were driven by the disappearance of taxa, more so for bacteria (days 17-95) than fungi (days 17-25; Table S7). Indeed, increased dominance by a few taxa, including the Firmicutes *Bacillus* at 25 days post-fire, *Paenibacillus* and *Domibacillus* (34-67 days), and the Proteobacteria *Noviherbaspirillum* (95 days), and the constant dominance of *Massilia* (34-286 days post-fire) resulted in the disappearance of early bacterial species, including RB41, *Conexibacter*, *Candidatus Udaeobacter*, and *Bacillus* over time (Fig. 5A; Table S7). In contrast, fungal succession was initiated by the rapid dominance of *Geminibasidium* and EMF genera that dominated the unburned communities, including *Inocybe*, *Cortinarius*, and *Tomentella* in the Basidiomycota and *Balsamia* in the Ascomycota and one previously rare Basidiomycete EMF genus *Mallocybe*, a subgenus of *Inocybe* (Fig. 5B, 6B). Interestingly, all species of EMF rapidly disappeared from the community at 25 days post-fire and were replaced by the constant dominance of *Pyronema*, *Penicillium*, and *Aspergillus* (Fig. 5B). However, as succession ensued, species appearance drove late-year successional turnover for bacteria (187-376 days; Fig. 5A) and mid-year successional turnover for fungi (67-187 days; Fig. 5B; Table S7). For bacteria, the appearance of two Bacteroidetes, *Pedobacter* and *Adhaeribacter*, and the Actinobacteria *Blastococcus* shaped late-year (days 187-376) bacterial succession (Fig. 5A). For fungi, mid-year succession (days 95-187) was driven by the appearance of filamentous fungal Ascomycetes in the Pyronemataceae and the Sordariaceae genus *Gelasinospora*, and Basidiomycota mushroom-forming taxa (Fig. 5B). For example, the previously dominant EMF genus *Inocybe* remained in the community but at a much lower abundance, declining with time since fire from 16% at 17 days to 0.1% at 376 days (Fig. 4D). In contrast, the pyrophilous Basidiomycete saprobes *Coprinellus* and *Tephrocybe*, which were rare in the unburned communities (<0.01% sequence abundance), appeared in the community later in the year (Fig. 5B). Late-year fungal succession (days 286-376) was driven by the increased dominance of *Tephrocybe* and *Aspergillus* and the Ascomycota *Rasamsonia*, and the decreased abundance of *Pyronema* (Fig. 5B).

**Table 1.** Pyrophilous bacterial and fungal taxa hypothesized position within Grime’s C-S-R succession theory (per traits at the genera level, based on current literature).

| Kingdom | Phylum | Genus | Grimmes Traits (CSR) | Trait description | Citation |
| --- | --- | --- | --- | --- | --- |
| Bacteria | Firmicute (Bacillota) | *Bacillus* | Survival | Aerobic, Heterotrophic, Antibiotic-producing, thermotolerant; endospore; reduces nitrogen; solubilizes phosphate | Grady et al., 2016; Kaur et al., 2018; Mandic-Mulec et al., 2015; J. et al., 2001; Kalayu, 2019; Slepecky and Hemphill in the Prokaryotes; Espinosa-de-los-Monteros et al., 2001 |
| Bacteria | Firmicute (Bacillota) | *Clostridiales* | Survival, Fast growth | Thermotolerant; endospore; trigger sporulation in response to stress, reduces nitrogen | Grady et al., 2016; Kaur et al., 2018; Mandic-Mulec et al., 2015; Paredes-Sabja et al., 2011; Kalayu, 2019 |
| Bacteria | Firmicute (Bacillota) | *Paenibacillus* | Survival | Aerobic, Heterotrophic, antibiotic producing (Bacteriocins), thermotolerant; endospore; fixes nitrogen | Grady et al., 2016; Kaur et al., 2018; Mandic-Mulec et al., 2015; Monciardini et al., 2003; Slepecky & Hemphill, 2006 |
| Bacteria | Actinobacteria (Actinomycetota) | *Conexibacter* | Competition | Aerobic, Heterotrophic, Antibiotic-producing, reduces nitrogen | Espinosa-de-los-Monteros et al., 2001; Kalayu, 2019 |
| Bacteria | Proteobacteria (Pseudomonadota) | *Massilia* | Fast Growth; Competition | Aerobic, fast reproduction, diverse ecology, exploit resources | Li et al., 2014; Toljander et al., 2005 |
| Bacteria | Proteobacteria (Pseudomonadota) | *Novihersparillum* | Fast Growth; Competition | Fast reproduction, exploit resources; likely PyOM degrader | Baldani et al., 2014; Grady et al., 2016; Woolet & Whitman, 2020 |
| Fungi | Basidiomycota | *Geminibasidium* | Survival | thermo- and xero-tolerant yeast | Alexopoulos & Mims, 1952 |
| Fungi | Ascomycota | *Pyronema* | Survival<br>Fast Growth | thermotolerant sclerotia; likely PyOM degrader, filamentous, | Moore, 1962; Fisher et al., 2022 |
| Fungi | Ascomycota | *Gelasinospora heterospora* | Survival | Requires heat or chemical treatment for germination, pigmented mycelia | Alexopoulos & Mims, 1952 |
| Fungi | Ascomycota | *Penicillium* | Fast Growth | copious asexual conidia, fast growth, filamentous | Dix & Webster, 1995; McGee et al., 2006; Crow, 1992 |
| Fungi | Ascomycota | *Aspergillus* | Fast Growth | copious asexual conidia, fast growth, filamentous | Dix & Webster, 1995; McGee et al., 2006; Crow, 1992 |
| Fungi | Basidiomycota | *Coprinellus* | Competition | Likely PyOM degrader, filamentous | Steindorff et al., 2021 |
| Fungi | Basidiomycota | *Tephrocybe anthrocophila* | Competition | Affinity for Ammonium or Nitrogen, filamentous | Suzuki, 2017; Legg, 1992 |

## 4. Discussion

Here, we present the highest resolution temporal sampling of post-fire microbiomes to date, enabling us to show for the first time that bacteria and fungi experience rapid succession driven by the dominance of pyrophilous taxa that traded off in abundance over time (H4). Although our study was in California chaparral, many of the dominant pyrophilous taxa identified here also dominate other burned environments, including Spanish shrublands (Pérez-Valera et al., 2018) and pine (Bruns et al., 2020; Reazin et al., 2016), spruce (Whitman et al., 2019), Eucalyptus (Ammitzboll et al., 2022; McMullan-Fisher et al., 2011), and redwood-tanoak forests (Enright et al., 2022), indicating the generality of our results to wildfire-affected ecosystems. We found that wildfire decreased bacterial and fungal biomass and richness, leading to community composition shifts that persisted the entire year (H1). Fungal guilds were differentially affected by fire and time, with AMF and saprobes experiencing the largest immediate fire impacts, but over time, EMF experienced the largest declines (H2). Moreover, microbial richness and biomass changes were driven by multiple abiotic interactions, including interactions between time since fire, precipitation, and soil burn severity (H3).

### 4.1 Wildfire decreased bacterial and fungal biomass and richness

Fire decreased soil bacterial and fungal biomass and richness, corroborating previous post-fire studies in Mediterranean shrublands (Pérez-Valera et al., 2018) and forests (Dooley & Treseder, 2012). We noted a larger fire effect on fungal biomass and richness relative to bacteria, consistent with previous research showing that bacteria are more resistant to fire than fungi (Certini et al., 2021; Glassman et al., 2021; Pourreza et al., 2014; Pressler et al., 2019). Although richness and biomass increased over time for both microbial groups, one year was insufficient for either group to recover to unburned levels, consistent with previous studies indicating that in shrublands, microbial biomass and richness recovery could take over two decades (Pérez-Valera et al., 2018). Interestingly, we observed a transient increase in bacterial richness directly post-fire, which may result from taxa such as the Actinobacteria genera *Soliribrobacter* and *Conexibacter* (Albuquerque & da Costa, 2014) that may be favored by the increase in soil pH observed here (Table S1) and in other studies (Neary et al., 1999) or by post-fire increases in nitrogen and phosphorus availability (Certini et al., 2021).

### 4.2 Wildfire impacts on fungal guilds

Fire-induced mortality was rapid for AMF and saprobes. Yet, EMF were most affected by fire over time, corroborating studies across ecosystems that point to the negative impact of fire on mycorrhizal richness, primarily driven by host mortality (Dove & Hart, 2017; Pulido-Chavez et al., 2021). Indeed, post-fire, all subplots were devoid of vegetation (0% vegetation cover) compared to 97% vegetation cover in unburned plots. However, we note that the rapid decline in AMF could be due to primer bias, as the ITS2 primer does not properly detect all AMF taxa (Lekberg et al., 2018). Moreover, we show that surviving ectomycorrhizal Basidiomycetes *Cortinarius*, *Inocybe*, and *Tomentella*, previously found in burned temperate pine forests (Owen et al., 2019; Pulido-Chavez et al., 2021) and pine-dominated Mediterranean systems (Gassibe et al., 2011; Hernández-Rodríguez et al., 2013) also dominated early chaparral fungal succession and remained in the community for 1-2 months. Since EMF largely disappeared after two months, this potentially answers the question of how long it takes mycorrhizal fungi to perish after their hosts’ death. Although it is possible that the EMF signal detected could be relic DNA (Carini et al., 2016), high soil temperatures typical of post-fire systems (Amacher et al., 2001; Neary et al., 1999) are likely to have degraded relic DNA (Sirois & Buckley, 2019; Torti et al., 2015), suggesting that these EMF could have survived the fire and clung to a dying host for at least two months. The survival of some EMF genera over others may be attributable to exploration type and specificity for current versus stored photosynthates (Gray & Kernaghan, 2020; Pena et al., 2010). For example, *Cortinarius*, a long/medium exploration type, which requires more carbon for maintenance, only survived for 17 days post-fire. In contrast, *Inocybe*, a short-contact type (Agerer, 2001; Koide et al., 2014), survived at very low abundance for the entire year, perhaps because it may be less carbon demanding or adapted to warming (Fernandez et al., 2017). Our results indicate that early EMF could have survived on their hosts’ stored photosynthates or the lower resources provided by the surviving and resprouting manzanita and chamise shrubs, while the vast majority of EMF experienced fire-induced mortality.

### 4.3 Pyrophilous fungi dominate the burned communities

Previously detected pyrophilous fungi across several ecosystems also dominated California chaparral, indicating that pyrophilous microbes are not biome-specific. Instead, these fungi may be activated by temperature thresholds reached during high-severity fires, which are common in chaparral and some Pinaceae forests (Agee, 1993; Keeley & Zedler, 2009; Neary et al., 1999). Specifically, our burned communities were dominated by Ascomycota in the genera *Pyronema*, *Penicillium*, and *Aspergillus*, similar to dominant post-fire fungal species in Mediterranean shrublands (Livne-Luzon et al., 2021), boreal spruce (Whitman et al., 2019), and montane pine forests (Bruns et al., 2020; Pulido-Chavez et al., 2021). Both *Pyronema* (*P. omphalodes*) and *Aspergillus* (*A. fumigatus*) are known pyrophilous fungi of California chaparral (Dunn et al., 1982), and we identified additional species including *P. domesticum*, *A. udagawae*, and *A. elsenburgensis*. These fungi are adapted to wildfire and produce fire-resistant structures, including dormant spores, sclerotia, and conidia, which are heat-activated (Gottlieb, 1950; Moore, 1962; Rhodes, 2006; Warcup & Baker, 1963). The rapid and efficient germination of *Aspergillus* (in the subgenus *Fumigati*, including *A. udagawae*) induced by high temperatures (Rhodes, 2006) and its ability to use various carbon and nitrogen sources, including ammonium and nitrate (Krappmann & Braus, 2005), may position *Aspergillus* to rapidly dominate post-fire. Furthermore, *Pyronema domesticum* can mineralize pyrogenic organic matter ((Fischer et al., 2021), an abundant substrate in post-fire environments. Moreover, *Geminibasidium*, a recently described thermotolerant Basidiomycete yeast (Nguyen et al., 2013), dominated at 17 days post-fire. While not typically described as pyrophilous, presumably because most pyrophilous fungi are described from mushrooms (McMullan-Fisher et al., 2011), two other studies have found *Geminibasidium* to increase post-fire in pine forests (Pulido-Chavez et al., 2021; Yang et al., 2020), suggesting that *Geminibasidium* is an underrepresented pyrophilous fungus. Together, these results indicate that dominant pyrophilous fungi are more widespread than expected and that their response to fire is likely due to fire adaptive traits and temperature thresholds.

### 4.4 Pyrophilous fungi drive fungal secondary succession

Although pyrophilous fungi are widespread across ecosystems, we lacked an understanding of how soon these pyrophilous fungi appear, their turnover rates, and if they change in abundance over time (Fox et al., 2022). Successional theory states that early successional stages are dominated by fast-growing or ruderal (R) organisms (Kinzig & Pacala, 2013) that tradeoff between stress tolerance (S) and competitive (C) life-history strategies over time (Grime, 1977; Zhang et al., 2018). Recent adaptations of Grime’s C-S-R to microbiomes suggest that pyrophilous microbes survive and thrive post-fire with traits analogous to plants, including post-fire resource acquisition (C), thermotolerant structures (S), and fast growth (R) (Enright et al., 2022; Whitman et al., 2019). Our data suggest that chaparral pyrophilous microbes fit into these trait categories (Table 1) and that tradeoffs among these traits might drive microbial succession.

Early post-fire succession was driven by surviving thermotolerant fungi (i.e., *Geminibasidium* and *Pyronema*), followed by fast-colonizers (i.e., *Penicillium* and *Aspergillus*), which were overtaken by competitive fungi capable of exploiting post-fire resources. (i.e., *Coprinellus* and *Tephrocybe*). Whereas *Geminibasidium* is thermo- and xero-tolerant (Nguyen et al., 2013), *Pyronema* produces thermotolerant sclerotia (Moore, 1962), enabling wildfire survival. Although both species are thermotolerant, differences in the morphological growth characteristics between the yeast *Geminibasidium* and the filamentous *Pyronema* may explain the tradeoff in dominance. For example, unicellular proliferation allowed *Geminibasidium* to dominate instantly, but the ability to forage for nutrients and rapidly increase surface colonization may allow the filamentous *Pyronema* to better dominate the open space. Interestingly, *Gelasinospora heterospora*, a fungus closely related to heat-activated *Neurospora crassa* (Dettman et al., 2001; Emerson, 1948), produces pigmented mycelia, and is known to require heat or chemical treatment for germination (Alexopoulous & Mims, 1952), was highly abundant at 25, 34, and 95 days post-fire, suggesting it is also pyrophilous. The decline of thermotolerators coincided with the dominance of fast-growing Ascomycete fungi in the genera *Penicillium* and *Aspergillus* (Dix & Webster, 1995; McGee et al., 2006), which both produce copious asexual conidia (Crow, 1992), allowing for rapid colonization of the open niche, analogous to early dominant post-fire plants, which are also often asexual (James, 1984). Lastly, like plants, competition appears to drive later successional dynamics (Tilman, 1990; Zhang et al., 2018), as suggested by the emergence of pyrophilous fungal decomposers, such as *Coprinellus* and *Tephrocybe* at 9-12 months, corroborating findings from Eucalyptus forests (Ammitzboll et al., 2022; McMullan-Fisher et al., 2011). Evidence suggests that *Coprinellus* can degrade aromatic hydrocarbons (Steindorff et al., 2021), supporting the idea that it is outcompeting earlier fungal taxa for this abundant and complex carbon source. Moreover, *Tephrocybe anthracophila* has a high affinity for Ammonium-Nitrogen (Legg, 1992; Suzuki, 2017), allowing access to this abundant post-fire resource. In conclusion, our results indicate that tradeoffs among fire-adapted traits analogous to those in plants might drive post-fire microbial succession.

### 4.5 Pyrophilous bacteria dominate and initiate bacterial secondary succession

Our high temporal sampling enabled us to demonstrate that bacteria experience rapid succession and community turnover initiated by possible aerobic, heterotrophic bacteria that form endospores and produce antibiotics, which may improve their competitive abilities in the post-fire environment. Moreover, we show that previously detected putative pyrophilous bacteria from Canadian (Whitman et al., 2019), Chinese boreal (Xiang et al., 2014), and California redwood-tanoak forests (Enright et al., 2022) and Spanish shrublands (Sáenz de Miera et al., 2020), also dominated California chaparral, suggesting that these pyrophilous bacteria might be widely distributed. Specifically, our burned communities favored taxa in the phyla Actinobacteria, Acidobacteria, Firmicutes, and Proteobacteria, similar to previous studies (Ammitzboll et al., 2022; Enright et al., 2022; Whitman et al., 2019; Xiang et al., 2014).

Bacterial successional dynamics differed from fungi in that the first few weeks were characterized by high diversity and a lack of dominance, mimicking the distributions observed in the unburned communities. In complex resource environments, such as those created by wildfire, niche complementarity is more important than competition because diverse communities can better exploit resources (Eisenhauer et al., 2013), further contributing to community stability. Our results support this idea, as the dominant taxa in our study are likely to use different resources. For example, members of the genus *Paenibacillus* can fix nitrogen (Monciardini et al., 2003; Slepecky & Hemphill, 2006), *Bacillus* and *Conexibacter* can reduce nitrogen, and *Bacillus* can further solubilize phosphate (Espinosa-de-los-Monteros et al., 2001; Kalayu, 2019). However, at 25 days, bacteria experienced a large compositional turnover, which allowed for the dominance of pyrophilous bacteria with fire-adapted traits. While tradeoffs in abundance were not as stark for bacteria as for fungi, as indicated by the constant dominance of the Proteobacteria *Massilia*, the following most abundant bacteria did experience tradeoffs in dominance against other bacterial genera, possibly driven by physiological traits, such as thermotolerance and fast colonization (Table 1).

Firmicutes contain some of the most resistant, thermotolerant endospore-producing bacteria (Grady et al., 2016; Kaur et al., 2018; Mandic-Mulec et al., 2015), including members of the genera *Bacillus*, *Paenibacillus*, and *Clostridiales* which decreased in abundance as fast-colonizing genera in the Phylum Proteobacteria that may be capable of exploiting post-fire substrates (e.g., *Novihersparillium* and *Massilia*) increased in dominance. Moreover, members of the Firmicute order Clostridiales can trigger sporulation in response to stress (Paredes-Sabja et al., 2011), potentially allowing for increased proliferation post-fire. While thermotolerance is crucial for fire survival, rapidly dominating the open niche and using post-fire resources may ensure post-fire growth. *Massilia* is a fast reproducing bacteria that associates with the rhizosphere (Li et al., 2014) and colonizes AMF hyphae (Iffis et al., 2014) and the roots of various plants (Ofek et al., 2012). Thus, the dominance of *Massilia* in our study and other post-fire environments (Enright et al., 2022; Whitman et al., 2019) suggests that its diverse ecology and rapid reproduction rate promote its proliferation within the soil (Toljander et al., 2005). Moreover, our results, coupled with current knowledge in the field, show that the continued dominance of *Massilia* and *Noviherbaspirillium* might be due to their ability to exploit post-fire resources. For example, the genera *Massilia* contains species that can degrade long-chain hydrocarbons (Ren et al., 2018) and reduce nitrates (Bailey et al., 2014). In contrast, *Noviherbaspirillium*, a mid-to late-year dominant bacterial genus, may degrade polycyclic aromatic hydrocarbon (Baldani et al., 2014; Grady et al., 2016; Woolet & Whitman, 2020). We conclude that although tradeoffs are not as distinctive for bacteria as they are fungi, potentially due to interaction among traits such as fast growth under high resources and resource acquisition under low resources, our results provide evidence that pyrophilous bacteria have mechanisms to survive the fire that may drive them to trade off in abundance from early to late dominant taxa over time.

### 4.6 Microbial successional dynamics differ for bacteria and fungi

Chaparral vegetation successional patterns result from the self-replacement of the dominant pre-fire species (i.e., auto-succession), meaning the pre-fire dominant species establish early post-fire due to resprouting individuals, and these species gradually dominate the system (Hanes, 1971). However, this was not the case for bacterial or fungal succession. Most post-fire dominant genera were rare or absent pre-fire, and these initial colonizers experienced directional change over time via species replacement. Our results are consistent with successional concepts of non-catastrophic disturbances, which show that species that survive a disturbance or recover quickly will dominate early successional dynamics but will inevitably be replaced by late-stage species via directional replacement mechanisms (Platt & Connell, 2003). Similar directional replacement patterns have been revealed after primary succession of fungi in a glacier foreland (Dong et al., 2016). Although a few genera dominated burned microbial communities, bacteria and fungi displayed different successional patterns. Bacterial burned communities exhibited lower synchrony and higher stability compared to unburned communities. The low synchrony indicates that stability may be conferred due to compensatory dynamics driven by tradeoffs in the abundance of cooccurring competitors (Tilman, 1999). Conversely, the lower stability in the unburned communities could be due to the increase in species synchrony reducing compensatory dynamics. In systems with high environmental variability, compensatory dynamics mediated by species asynchrony often contribute to stability (Yachi & Loreau, 1999). In contrast, burned fungal communities displayed lower community stability compared to unburned communities, while synchrony was similarly low in both. This lower stability may indicate that there was an overall increase in the variation of fungal taxa potentially due to the large loss of species observed in the fungal community (Tilman et al., 2014), resulting in priority effects and alternate communities with different dominant genera (Connell & Slatyer, 1977; Fukami, 2015). However, further research is needed to explore these microbial successional dynamics, as microbial communities are a vital driver of the post-fire ecosystem recovery.

### 4.7 Precipitation and burn severity differentially affected bacterial and fungal communities

Although fire was the most influential driver of microbial change, we found that interactions between time and precipitation also drove successional dynamics by differentially affecting microbial biomass and richness. Precipitation interacted with time since fire resulting in a negative correlation with bacterial richness and a positive correlation with fungal richness, with bacterial richness declining but fungal richness increasing after rain events. At least one study that performed a three-year precipitation manipulation experiment showed that increased precipitation decreased bacterial richness (Yang et al., 2021). In contrast to richness, precipitation interacted with fire to positively affect bacterial biomass, while fungal biomass remained unaffected by rainfall. The distinct responses to precipitation could be due to microbial physiological and morphological differences (Blazewicz et al., 2014; Placella et al., 2012; Selbmann et al., 2013). For example, fungi have high morphological plasticity, including the ability to alternate growth forms based on environmental conditions, such as hyphal growth in low nutrient conditions versus unicellular form in rich nutrient conditions (Selbmann et al., 2013), which may allow fungi to survive small changes in moisture content, allowing for continued growth over time. Moreover, fungi can also activate or enhance resistance to drought (Evans & Wallenstein, 2012; Guhr & Kircher, 2020), allowing them to maintain a constant reproduction rate regardless of the external environmental conditions, thus explaining the positive impact on fungal richness. In contrast, bacteria can experience cell death due to rapid wet-up, while other bacterial groups enter a dormant state, inhibiting growth (Schimel, 2018). Thus, to be evolutionarily successful, some bacteria may respond rapidly to favorable conditions (e.g., rain) (Leizeaga et al., 2022) and rapidly replicate to outcompete other microbes for the pulse of nutrients released during the rain event (Homyak et al., 2014), thus potentially explaining the peaks in richness observed after significant rainfall at day 131.

Previous research shows that ash depth correlates with fire intensity (Rice, 1993) and severity (Bodí et al., 2014; Parson et al., 2010). Here, we measured ash depth before the first rains, or significant wind events could remove ash from the site, and we found that ash depth can serve as a proxy of soil burn severity on a scale relevant to soil microbes. Unlike the BAER maps, which classified all plots as moderate burn severity, we identified large variations in soil burn severity across our subplots (Fig. S7). It is well known that wildfires result in a heterogeneous landscape (Jain et al., 2008); thus, the discrepancy in soil burn severity in the BAER maps could be due to the lack of ground verification. Since our severity effects were similar to those found in other studies, we feel confident that ash depth can serve as a proxy for soil burn severity. We found that severity negatively affected bacterial and fungal richness but not biomass. Similar to studies in boreal (Whitman et al., 2019), oak (Pourreza et al., 2014), and Mediterranean forests (Lucas-Borja et al., 2019), bacterial and fungal richness decreased with soil burn severity. However, contrary to a meta-analysis (Dooley & Treseder, 2012), neither bacterial nor fungal biomass was directly impacted by soil burn severity, probably due to microbial adaptations to the natural high severity fire regime of chaparral. However, since we observed an interaction effect of time since fire and severity on microbial richness, it is possible that previously observed severity effects, derived from chronosequence studies where space is substituted for time and which, on average, occur two years post-fire (Pressler et al., 2019), may represent an interaction effect of time and severity and not a direct effect of soil burn severity.

## 5. Conclusion

Our high-resolution temporal sampling allowed us to detect rapid secondary succession in bacterial and fungal communities and marked tradeoffs in abundance among pyrophilous microbes over time. We found that the same pyrophilous bacteria and fungi that respond to fires in temperate and boreal forests and Mediterranean shrublands also dominate in California chaparral, allowing us to generalize knowledge of post-fire microbiomes to dryland habitats that are rapidly increasing with climate change (Feng & Fu, 2013). While pyrophilous microbes have been described in other systems, their turnover rates were unknown due to limited post-fire sampling (Fox et al., 2022). We hypothesize that tradeoffs among fire-adapted traits, with thermotolerators dominating first, followed by fast colonizers, and finally by competitors capable of capitalizing on post-fire resource acquisition, drive post-fire microbial successional dynamics. It appears likely that pyrophilous microbes share many analogous traits to plants, enabling us to adapt successional theory developed for plants to microbes while acknowledging that bacteria and fungi differ in how they respond to fire according to their physiology. We conclude that post-fire bacteria and fungi experience rapid successional changes after fire, suggesting that these dynamics can help increase our ability to predict if and how ecosystems recover after a wildfire, an increasingly widespread disturbance with climate change.

## Acknowledgments

We thank the Cleveland National Forest and the Trabuco Ranger District, including District Ranger Darrel Vance and Emily Fudge, Jeffrey Heys, Lauren Quon, Jacob Rodriguez, and Victoria Stempniewicz, for their guidance and help with permitting and site selection. We thank Judy A. Chung for her help with fieldwork and molecular work and Aral C. Greene, Dylan Enright, and Sameer S. Saroa for fieldwork assistance. We thank Amelia Nelson, Elizah Stephens, Kaleigh A. Russell, and anonymous reviewers for their helpful comments on the manuscript. This work was supported by UC Riverside, the BLM JFSP Award #012641-002 and Shipley Skinner Award to MFPC and SIG, the USDA-NIFA Award 2022-67014-36675 to SIG and PMH, and the DOE BER Award DE-SC0023127 to SIG and PMH.

## Data Accessibility

Raw sequence reads were submitted to the National Center for Biotechnology Information Sequence Read Archive under BioProject accession number PRJNA761539. All statistical codes are available on GitHub https://github.com/pulidofabs/SecondarySuccession-Chaparral.

## Benefit-Sharing

Nagoya Protocol is not applicable as there is no genetic material to report. However, benefits from this research ensue from the sharing of our data and results on public databases as described above.

## Author’s contribution

SIG conceived of the study and acquired permits; LL, SIG, and PMH came up with the experimental design; SIG, LL, PMH, MFPC, and JWJR performed fieldwork; MFPC and JWJR performed all molecular work; MFPC performed all bioinformatics, statistical analysis, made figures, and wrote the first draft; all authors contributed to writing and editing of the manuscript.

