## Supplementarty Files for "Rapid bacterial and fungal successional dynamics in first year after Chaparral wildfire"

MOLECULAR ECOLOGY

**Supplemental Information for:**

### Bacterial and fungal communities experience rapid secondary succession du ring the first year following a wildfire in California chaparral

M. Fabiola Pulido-Chavez, James W. J. Randolph, Cassandra Zalman, Loralee Larios, Peter M. Homyak, and Sydney I. Glassman

#### Table of Contents:


| | |
|---|---|
| **Figure S1.** Alpha diversity metrics for bacteria and fungi between treatments | Page 1 |
| **Figure S2.** Change in average bacterial and fungal biomass between treatments | Page 2 |
| **Figure S3.** Change in average richness for the fungal guilds | Page 3 |
| **Figure S4.** Changes in bacterial and fungal species richness over time | Page 4 |
| **Figure S5.** NMDS plots for bacterial community composition at each time points | Page 5 |
| **Figure S6.** NMDS plots for fungal community composition at each time points | Page 6 |
| **Table S1.** Holy Fire Site-specific characteristics for the nine sampling plots | Page 7 |
| **Table S2.** Descriptive statistic and percent change in biomass and species richness | Page 8 |
| **Table S3.** Negative binomial generalized mixed effect models for bacteria and fungi | Page 9 |
| **Table S4.** Negative binomial generalized mixed effect models for the fungal guilds | Page 1[illegible] |
| Table S5. Permanova of bacterial and fungal community composition and treatment | Page 1[illegible] |
| Table S6. Measures of successional dynamics for bacterial and fungal communities | Page 1[illegible] |

**Figure S1.** Comparison of alpha diversity metrics for bacteria and fungi between burned (brown) and unburned (blue-green) plots for A) observed species richness (ASVs), B) Simpson, C) Chao1, D) ACE, E) Inverse Simpson (dominance), and E) Shannon. Significance based on negative binomial regression with plots and time since fire as random effects for all alpha metrics except for bacteria inverse Simpson which was based on a generalized mixed effect model. Percent value represents the percent change in alpha diversity from the unburned to burned communities, where the negative value represents a decrease in alpha diversity.

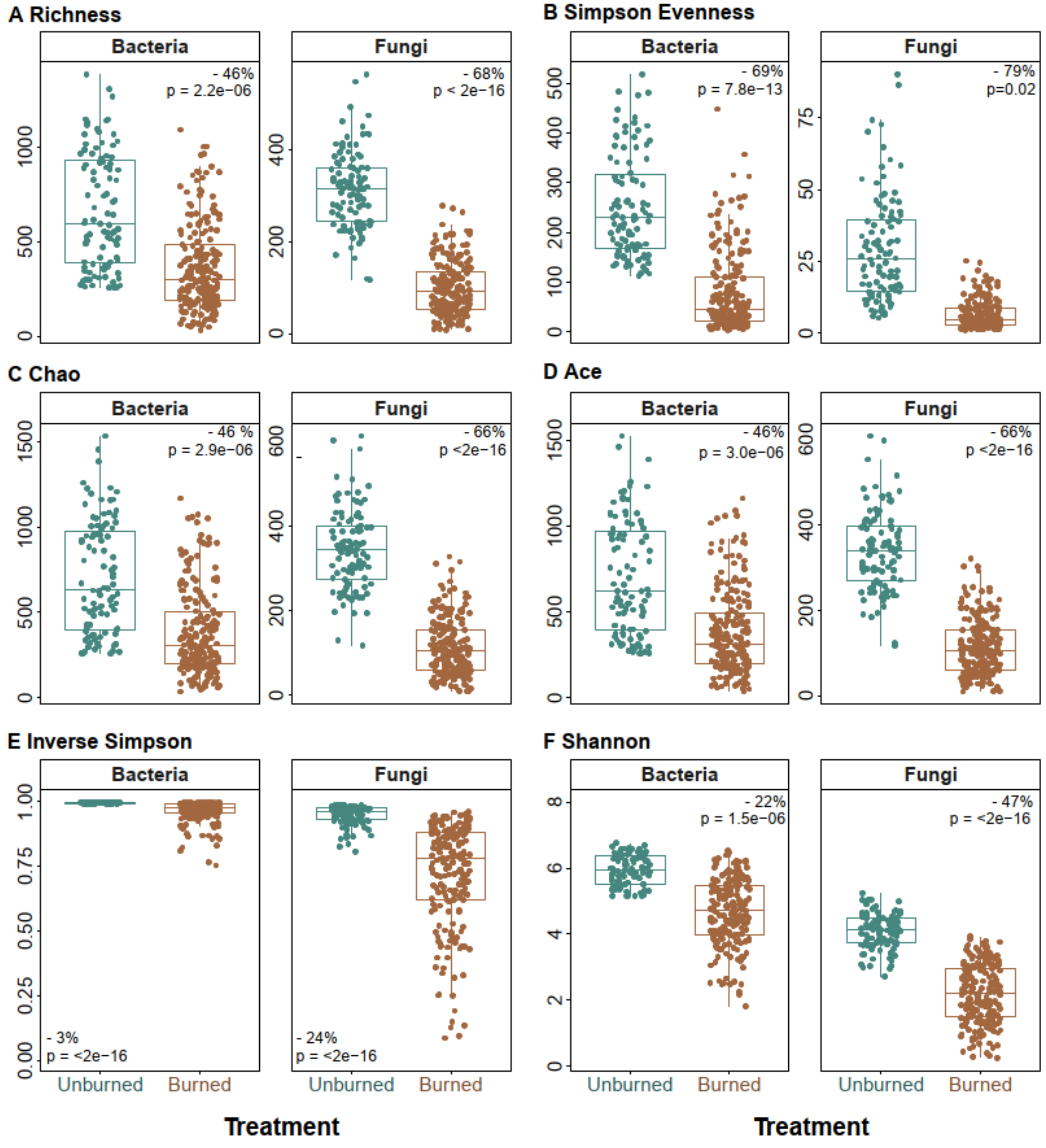

**Figure S2**. Change in average A) bacterial and B) fungal biomass between the burned (brown) and unburned (blue-green) communities across all time points. Boxes represent the 25th and 75th quartiles, and the horizontal line is the median of the data—significance based on negative binomial regressions with plot and time since fire as random effects. Percent value represents the percent change in biomass from the unburned to burned communities, where the negative value represents a decrease in biomass.

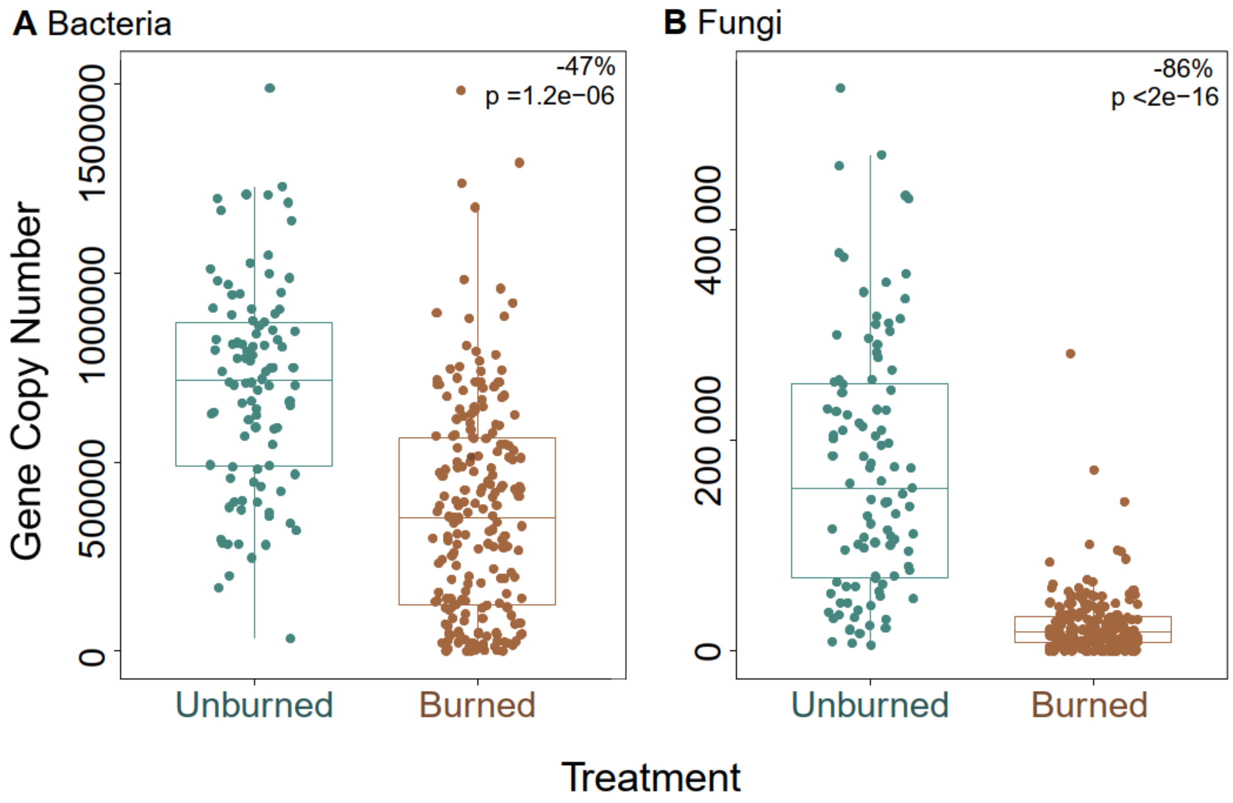

**Figure S3.** Change in average richness for A) arbuscular mycorrhizal fungi (AMF), B) ectomycorrhizal fungi (EMF), C) pathogenic, and D) saprobic fungi between burned (brown) and unburned (blue-green) plots across all time points. Boxes represent the 25th and 75th quartiles, and the horizontal line is the median of the data. Significance based on negative binomial regressions with plot, subplot, and time since fire as random effects. Percent value represents the percent change in species richness from the unburned to burned communities, where the negative value represents a decrease in richness.

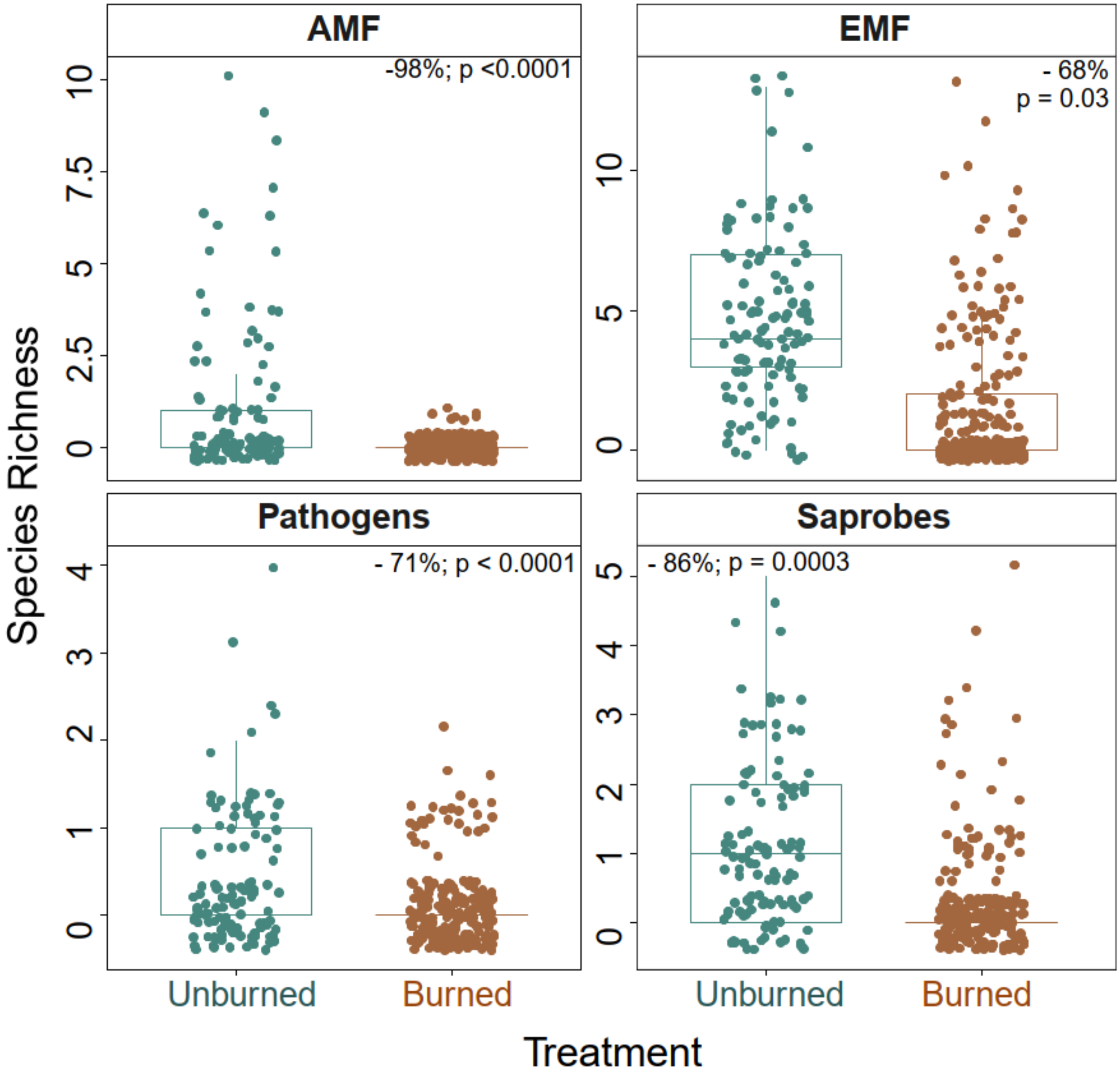

**Figure S4.** Change of species richness in burned (brown) versus unburned (blue-green) plots at each of the 9-time points for all four fungal guilds (arbuscular mycorrhizal fungi (AMF); ectomycorrhizal fungi (EMF), pathogens, and saprobes). Points represent the mean, and bars represent the standard error of the mean.

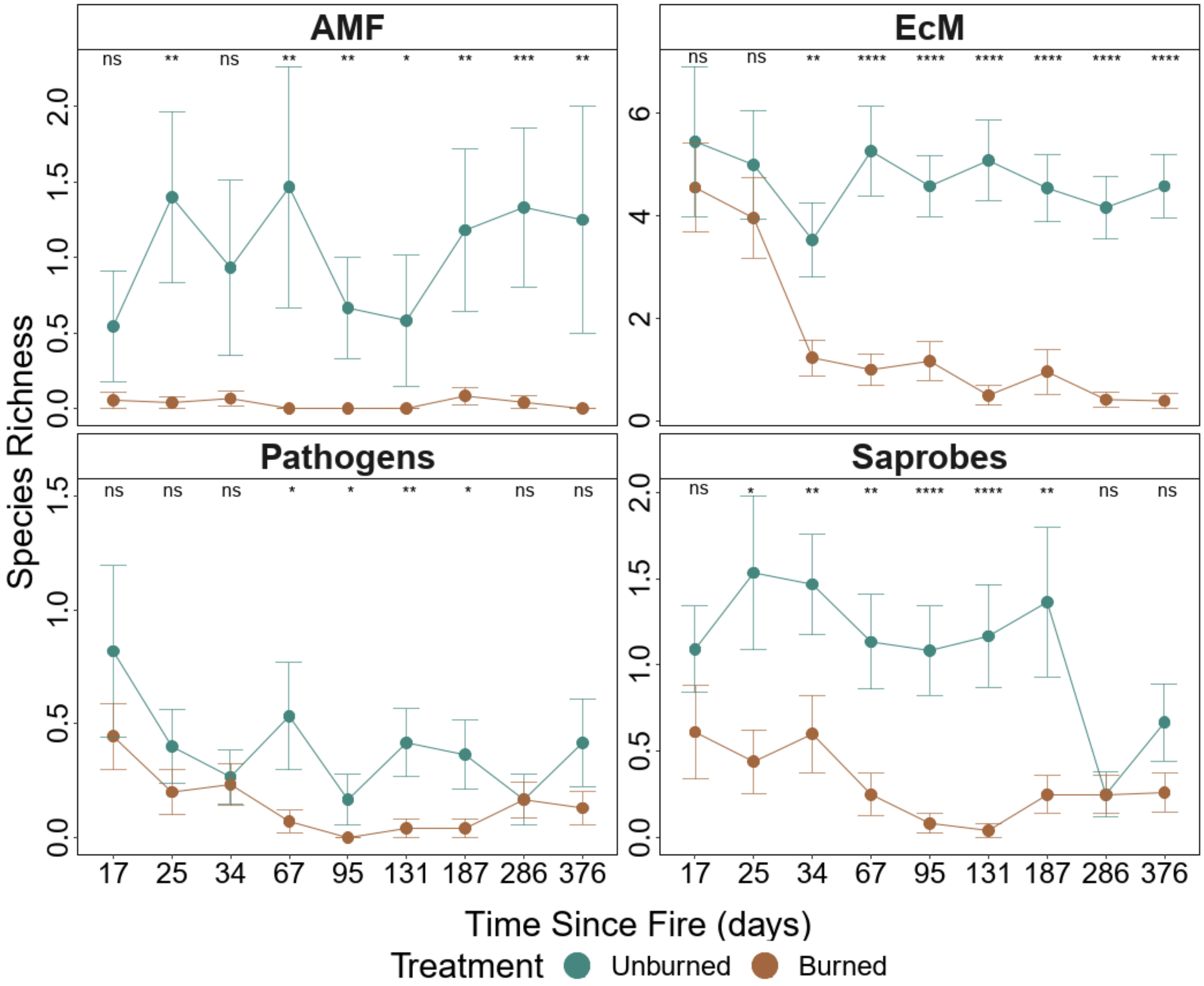

**Figure S5.** NMDS plots for bacterial community composition in burned (brown) versus unburned (blue-green) plots at each of the 9-time points with $R^2$, significance, and stress (S) based on ADONIS. NMDS is based on the Bray-Curtis dissimilarity matrix on 3-dimensions.

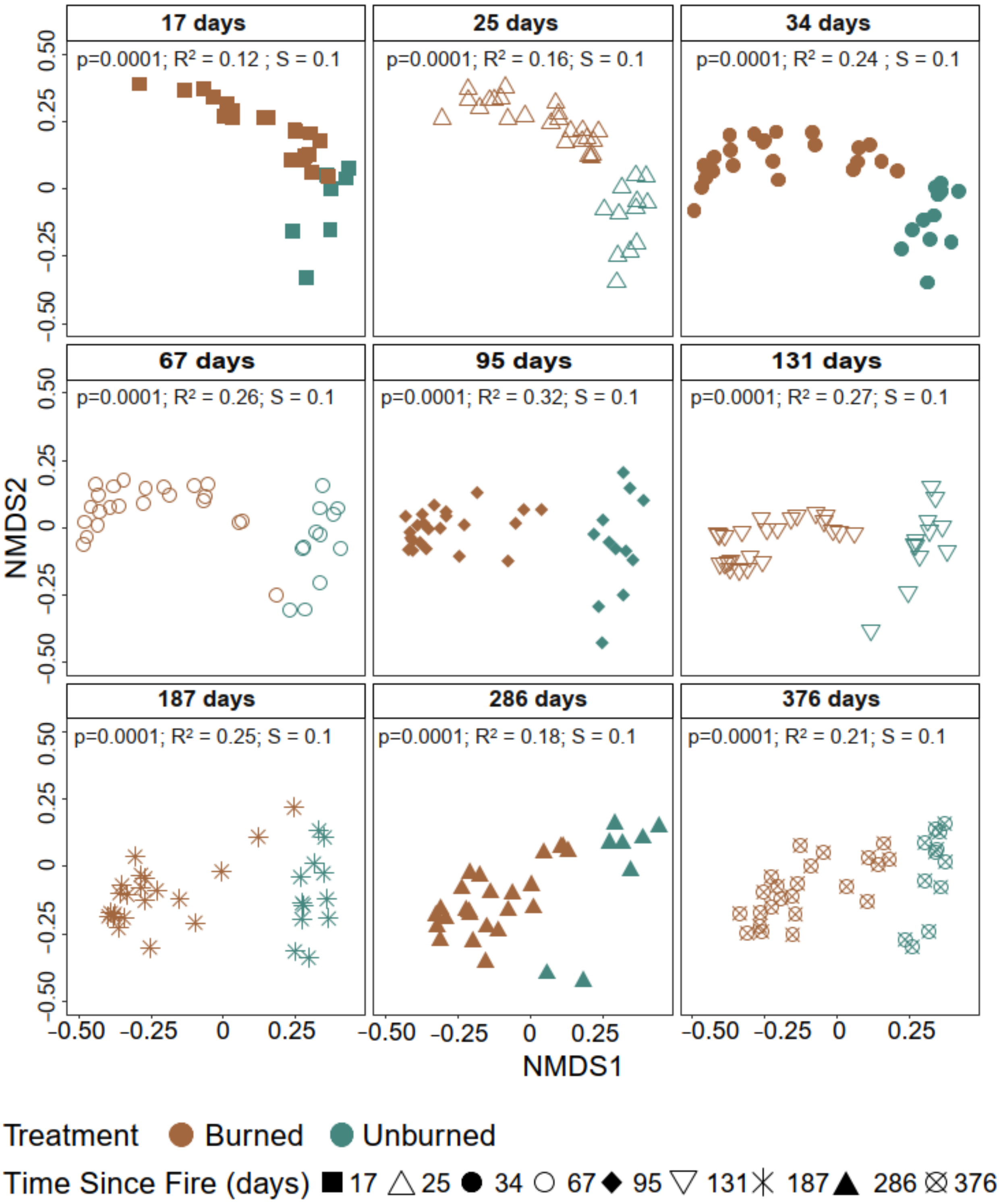

**Figure S6.** NMDS plots for fungal community composition in burned (brown) versus unburned (blue-green) plots at each of the 9-time points with $R^2$, significance, and stress (S) based on ADONIS. NMDS is based on the Bray-Curtis dissimilarity matrix on 3-dimensions.

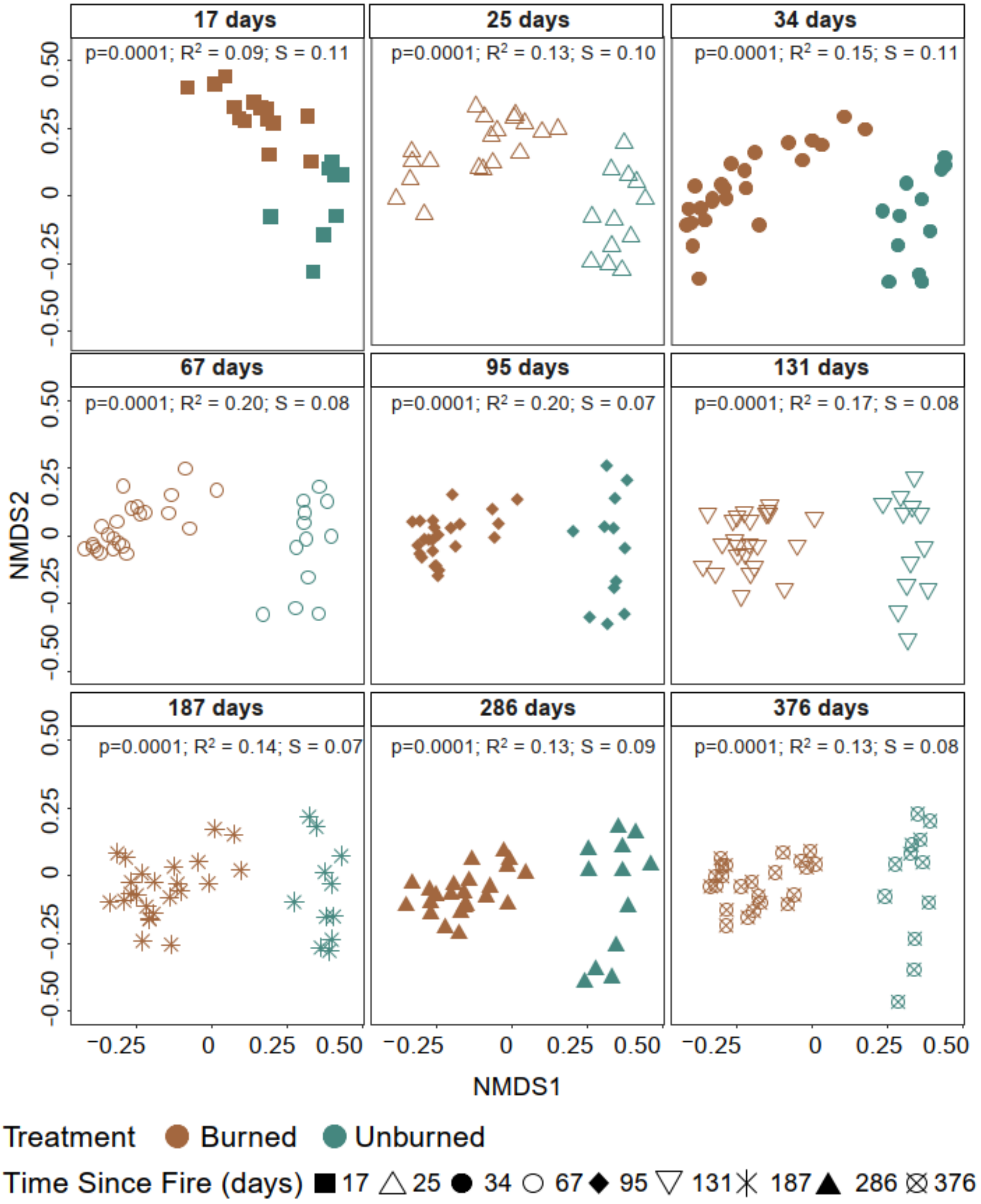

**Table S1.** Holy Fire Site-specific characteristics for the nine sampling plots (6 burned, 3 unburned) located within the 2018 Holy Fire.

| Site ID | Treatment | Latitude | Longitude | Elevation | Soil pH | | Soil Taxonomic Class |
|---|---|---|---|---|---|---|---|
| | | | | | 25 days | 376 days | |
| CNF01 | Burned | 33.6901 | -117.463 | 1228 | 7.25 | 6.33 | Cieneba series; Loamy, mixed, superactive, nonacid, thermic, shallow Typic Xerorthents (Entisols) |
| CNF02 | Burned | 33.69537 | -117.471 | 1260 | 6.71 | 6.74 | |
| CNF03 | Burned | 33.69326 | -117.467 | 1237 | 6.79 | 7.1 | |
| CNF04 | Burned | 33.68456 | -117.457 | 1260 | 7.11 | 6.84 | Friant series; Loamy, mixed, superactive, thermic Lithic Haploxerolls (Mollisols) |
| CNF05 | Burned | 33.67809 | -117.457 | 1195 | 6.94 | 7.17 | |
| CNF06 | Burned | 33.67168 | -117.459 | 1285 | 6.92 | 7.48 | |
| CNF07 | Unburned | 33.67135 | -117.459 | 1283 | 6.1 | 6.85 | |
| CNF08 | Unburned | 33.66813 | -117.456 | 1250 | 6.12 | 6.88 | |
| CNF09 | Unburned | 33.6678 | -117.455 | 1240 | 6.18 | 6.18 | |

**Table S2.** Descriptive statistic and percent change in biomass and species richness between treatment (unburned (burned)) and time since fire for bacteria and fungi. The mean copy number is based on the 16S rRNA for bacteria and 18S rRNA for fungi.

| | | Biomass | | Richness | |
|---|---|---|---|---|---|
| | **Time since fire (days)** | **Mean Copy Num.** | **% Change** | **Mean ASVs** | **% Change** |
| **Bacteria** | 17 | 608221(99082) | -84 | 462(604) | 31 |
| | 25 | 587923(91326) | -85 | 688(486) | -29 |
| | 34 | 631138(267965) | -58 | 962(313) | -67 |
| | 67 | 734587(330466) | -55 | 621(250) | -60 |
| | 95 | 824341(613725) | -26 | 302(180) | -41 |
| | 131 | 692448(419958) | -39 | 972(378) | -61 |
| | 187 | 728258(603165) | -17 | 933(244) | -74 |
| | 286 | 807788(424704) | -47 | 368(411) | 12 |
| | 376 | 707535(401262) | -43 | 533(411) | -23 |
| **Fungi** | 17 | 209062(5865) | -97 | 292(161) | -45 |
| | 25 | 188925(7669) | -96 | 369(117) | -68 |
| | 34 | 225263(26888) | -88 | 305(77) | -75 |
| | 67 | 177830(36099) | -80 | 328(60) | -82 |
| | 95 | 135010(25283) | -81 | 335(85) | -75 |
| | 131 | 108756(12465) | -89 | 312(93) | -70 |
| | 187 | 133283(27691) | -79 | 344(118) | -66 |
| | 286 | 217373(41395) | -81 | 268(119) | -55 |
| | 376 | 182563 (37027) | -80 | 257(99) | -61 |

**Table S3.** Model summary results of the effect of treatment (burned vs. unburned), time since fire (TSF in days), precipitation (mm), and soil burn severity measured as ash depth (cm) at day 17 on bacterial and fungal biomass and richness. Significance is based on the negative binomial generalized mixed effect models with plot, subplot, and time since fire as the random effect for richness and plot and time since fire as random effect for biomass—significance at $p<0.05$ (bold).

| | Bacteria | | | Fungi | | |
|---|---|---|---|---|---|---|
| | **Est.** | **z value** | **P value** | **Est.** | **z value** | **P value** |
| **Biomass** | | | | | | |
| (Intercept) | 13.45 | 69.65 | **<0.0001** | 12.02 | 55.45 | **<0.0001** |
| Treatment (Burned) | -0.82 | -3.85 | **0.0001** | -2.13 | -8.01 | **<0.0001** |
| TSF | 0.10 | 0.45 | 0.65 | 0.08 | 0.56 | 0.57 |
| Precipitation | 0.06 | 0.43 | 0.67 | - | - | - |
| Soil burn severity (ash depth) | -0.01 | -0.49 | 0.62 | 0.00 | 0.01 | 0.99 |
| Treatment (Burned):Precipitation | 0.30 | 3.10 | **0.002** | - | - | - |
| TSF x Precipitation | -0.07 | -0.29 | 0.77 | - | - | - |
| TSF x Soil burn severity (ash depth) | 0.06 | 3.41 | **0.001** | 0.06 | 2.95 | **0.003** |
| **Random Effects** | | | | | | |
| Variance/Std.Dev | $0.08/0.29_{Plot;}$ $0.10/0.32_{TSF}$ | | | $0.09/0.30_{Plot;}$ $0.12/0.34_{TSF}$ | | |
| Mar. $R^2$ / Cond. $R^2$ | 0.30/0.46 | | | 0.48/0.58 | | |
| **Richness** | | | | | | |
| (Intercept) | 6.46 | 47.27 | **<0.0001** | 5.8 | 54.69 | **<0.0001** |
| Treatment (Burned) | -0.37 | -2.56 | **0.01** | -0.9 | -5.76 | **<0.0001** |
| TSF | -0.09 | -0.84 | 0.4 | 0.19 | 2.13 | **0.03** |
| Precipitation | 0.15 | 1.38 | 0.17 | -0.01 | -0.36 | 0.72 |
| Soil burn severity (ash depth) | -0.07 | -3.83 | **0.0001** | -0.07 | -3.48 | **0.001** |
| TSF x Precipitation | -0.27 | -4.64 | **<0.0001** | 0.34 | 3.272 | **0.001** |
| TSF x Soil burn severity (ash depth) | 0.02 | 2.38 | **0.02** | 0.03 | 3.82 | **0.0001** |
| **Random Effects** | | | | | | |
| Variance/Std.Dev | $0.02/0.13_{Plot}$ $0.02/0.14_{Subplot}$ $0.08/0.28_{TSF}$ | | | $0.02/0.12_{Plot}$ $0.04/0.20_{Subplot}$ $0.01/0.08_{TSF}$ | | |
| Mar $R^2$ / Cond. $R^2$ | 0.32/0.57 | | | 0.63/0.74 | | |

**Table S4.** Model summary results of the effect of treatment (burned vs. unburned), time since fire (TSF in days), precipitation (mm), and ash depth (cm) on arbuscular fungi (AMF), ectomycorrhizal fungi (EMF), Saprobes and Pathogens. Significance based on the negative binomial generalized mixed effect models with plot, subplot, and time since fire as random effect—significance at $p<0.05$ (bold).

| | **AMF** | | | **EMF** | | |
|---|---|---|---|---|---|---|
| | **Est.** | **z value** | **P value** | **Est.** | **z value** | **P value** |
| (Intercept) | -0.98 | -1.71 | 0.09 | 1.52 | 4.57 | **<0.0001** |
| Treatment (Burned) | -4.10 | -4.34 | **<0.0001** | -1.09 | -2.18 | **0.03** |
| TSF | - | - | - | -0.09 | -1.25 | 0.21 |
| Soil burn severity (ash depth) | - | - | - | -0.19 | -2.77 | **0.01** |
| Precipitation | - | - | - | 0.00 | -0.06 | 0.95 |
| Treatment (Burned) x Precipitation | - | - | - | -0.37 | -3.92 | **<0.0001** |
| Treatment (Burned) x TSF | - | - | - | -0.68 | -7.04 | **<0.0001** |
| **Random Effects** | | | | | | |
| Variance/Std.Dev | 0.23/0.48$_{Plot}$ 0.45/0.67$_{Subplot}$ 0.03/0.17$_{TSF}$ | | | 6.3e$^{-10}$/2.5e$^{-05}$$_{Plot}$ 3.06/1.75$_{Subplot}$ 0.06/0.25$_{TSF}$ | | |
| Mar $R^2$ / Cond. $R^2$ | 0.55/0.81 | | | 0.20/0.37 | | |

| | **Saprobes** | | | **Pathogens** | | |
|---|---|---|---|---|---|---|
| | **Est.** | **z value** | **P value** | **Est.** | **z value** | **P value** |
| (Intercept) | -0.06 | -0.35 | 0.73 | -1.28 | -4.83 | **<0.0001** |
| Treatment (Burned) | -1.96 | -7.37 | **<0.0001** | -1.48 | -3.63 | **0.0003** |
| TSF | -0.36 | -3.07 | **0.002** | - | - | - |
| Precipitation | - | - | - | 0.09 | 0.52 | 0.60 |
| Treatment (Burned) x Precipitation | - | - | - | -0.88 | -2.16 | **0.03** |
| Treatment (Burned) x TSF | 0.68 | 3.42 | **0.001** | - | - | - |
| **Random Effects** | | | | | | |
| Variance/Std.Dev | 1.01e$^{-11}$/3.2e$^{-06}$$_{Plot}$ 0.18/0.42$_{Subplot}$ 0.01/0.08$_{TSF}$ | | | 7.2e$^{-12}$/2.7e$^{-06}$$_{Plot}$ 0.34/0.58$_{Subplot}$ 3.9e$^{-13}$/6.3e$^{-07}$$_{TSF}$ | | |
| Mar $R^2$ / Cond. $R^2$ | 0.21/0.25 | | | 0.11/0.16 | | |

**Table S5.** Permutational multivariate analysis of variance (PERMANOVA) of bacterial and fungal community composition and the effects of treatment (burned vs. unburned), time since fire (TSF) in days, and their respective interactions. Significance $p<0.05$ (bold).

| | Variable | Sum of Sqs. | R2 | F | P value |
|---|---|---|---|---|---|
| | Treatment | 14.36 | 0.13 | 47.18 | **0.0001** |
| Bacteria | TSF days | 4.44 | 0.04 | 14.58 | **0.0001** |
| | Treatment: TSF days | 2.29 | 0.02 | 7.51 | **0.0001** |
| | Treatment | 13.85 | 0.11 | 37.67 | **0.0001** |
| Fungi | TSF days | 1.77 | 0.01 | 4.81 | **0.0001** |
| | Treatment: TSF days | 1.13 | 0.01 | 3.06 | **0.0003** |

**Table S6.** Measures of successional dynamics for bacterial and fungal communities between treatments (burned vs. unburned) where the unburned values are inside the parenthesis. Turnover rates (proportion of species that differ between time points), appearance (relative species appearance between time points) and disappearance (relative species disappearance between time points), rates of change (rate of directional change in community composition over time), Stability (of total species abundance as a measure of equilibrium) and Synchrony (a measure of whether abundance fluctuations are homo- or heterogeneous over time). Higher values represent a higher rate for each category.

| Microbe | TSF days | Turnover Rate | Appearance | Disappearance | Rate of change | Stability | Synchrony |
|---|---|---|---|---|---|---|---|
| Bacteria | 25 | 0.59 (0.54) | 0.25 (0.40) | 0.34 (0.14) | 0.16 (0.12) | 8.35 (5.36) | 0.03 (0.22) |
| | 34 | 0.51 (0.50) | 0.15 (0.21) | 0.36 (0.29) | | | |
| | 67 | 0.45 (0.43) | 0.21 (0.27) | 0.24 (0.16) | | | |
| | 95 | 0.50 (0.56) | 0.15 (0.14) | 0.35 (0.42) | | | |
| | 131 | 0.37 (0.40) | 0.19 (0.16) | 0.19 (0.24) | | | |
| | 187 | 0.50 (0.58) | 0.31 (0.21) | 0.19 (0.38) | | | |
| | 286 | 0.50 (0.64) | 0.28 (0.32) | 0.22 (0.32) | | | |
| | 376 | 0.50 (0.35) | 0.26 (0.25) | 0.24 (0.10) | | | |
| Fungi | 25 | 0.61 (0.50) | 0.26 (0.37) | 0.34 (0.13) | 0.49 (0.08) | 6.42 (8.58) | 0.04 (0.05) |
| | 34 | 0.52 (0.43) | 0.14 (0.30) | 0.38 (0.13) | | | |
| | 67 | 0.52 (0.53) | 0.28 (0.28) | 0.24 (0.25) | | | |
| | 95 | 0.42 (0.47) | 0.27 (0.25) | 0.15 (0.23) | | | |
| | 131 | 0.52 (0.57) | 0.29 (0.27) | 0.23 (0.30) | | | |
| | 187 | 0.55 (0.53) | 0.40 (0.24) | 0.15 (0.29) | | | |
| | 286 | 0.53 (0.53) | 0.15 (0.24) | 0.38 (0.29) | | | |
| | 376 | 0.42 (0.58) | 0.19 (0.31) | 0.23 (0.27) | | | |
